# Scalable *in situ* single-cell profiling by electrophoretic capture of mRNA

**DOI:** 10.1101/2022.01.12.476082

**Authors:** Lars E. Borm, Alejandro Mossi Albiach, Camiel C.A. Mannens, Jokubas Janusauskas, Ceren Özgün, David Fernández-García, Rebecca Hodge, Ed S. Lein, Simone Codeluppi, Sten Linnarsson

**Author notes:** Corresponding author. Correspondence to: L.E.B. | @larsborm, mS.L. | @slinnarsson | linnarssonlab.org.

## Abstract

Methods to spatially profile the transcriptome are dominated by a trade-off between resolution and throughput. Here, we developed a method named EEL FISH that can rapidly process large tissue samples without compromising spatial resolution. By electrophoretically transferring RNA from a tissue section onto a capture surface, EEL speeds up data acquisition by reducing the amount of imaging needed, while ensuring that RNA molecules move straight down towards the surface, preserving single-cell resolution. We applied EEL on eight entire sagittal sections of the mouse brain and measured the expression patterns of up to 440 genes to reveal complex tissue organisation. Moreover, EEL enabled the study of challenging human samples by removing autofluorescent lipofuscin, so that we could study the spatial transcriptome of the human visual cortex. We provide full hardware specification, all protocols and complete software for instrument control, image processing, data analysis and visualization.

## Introduction

The brain is a highly structured organ that comprises a vast diversity of cell types including many subtypes of excitatory and inhibitory neurons, astrocytes, oligodendrocytes, vascular and immune cells^1^. Cell numbers vary by cell type, from a few hundred cells (e.g. hypothalamus *Pmch+* neurons) to billions (e.g. cerebellar granule neurons) and are organized in intricate spatial patterns on scales ranging from micrometers to centimeters. Studying the organization and spatial relationships between all these different cell types by microscopy is challenging because the diversity vastly outnumbers the handful of colours that can be simultaneously resolved. However, recent advances in multiplexing have made it possible to elucidate the complex structure of the brain *in situ* at higher resolution in health and disease^2–6^.

These newer spatial methods use either sequencing or microscopy to detect mRNA in space^7,8^. Sequencing-based methods transfer the mRNA of a tissue section onto a surface where transcripts are hybridized or ligated to spots of spatially barcoded oligonucleotides, which are then collected and resolved by sequencing^9–15^. These methods can measure the entire transcriptome using existing sequencing pipelines and can scale to large areas and high throughput by parallel sample processing. Nevertheless, they are limited by low capture efficiency, gaps between capture spots and high sequencing costs. Furthermore, spot-based spatial data is difficult to segment into true single-cell data and is commonly analyzed in terms of spots, not cells.

Microscopy-based methods rely on single molecule Fluorescence *in situ* Hybridization (smFISH) or *in situ* sequencing to directly visualize RNA transcripts in tissue^2,5,6,16–21^. To be able to map many transcripts and overcome the limited color space resolvable by fluorescent microscopes, some smFISH methods use a barcoded approach in which different gene transcripts are encoded by a binary barcode that is detected by multiple cycles of labelling, imaging and stripping. Barcoding enables the detection of as many as 10,000 different RNA species^4,22^. Nevertheless, the analysis of large samples is time-consuming because of repeated imaging at high magnification using high-numerical aperture microscope objectives. The small field of view (FOV) and shallow depth of field of such objectives necessitate the imaging of many XY tiles and a large Z-stack, so that surveying only a few mm^2^ can take days^3^ to weeks^2^.

One way to increase the imaging speed is to make the signal detectable by low power microscope objectives characterised by a large FOV through signal amplification but sacrificing quantitative sensitivity^5,6,21^. However, the reduced resolution of low magnification objectives causes an increase in optical crowding, in which signals from multiple RNA molecules cannot be separated, therefore preventing high multiplexing (Extended Data Fig. 1a,b).

Here, we describe Enhanced ELectric (EEL) FISH, an smFISH-based method that combines high multiplexing with large area imaging at high resolution. By electrophoretically transferring RNA onto a glass surface and removing the tissue, time-consuming imaging in the Z axis can be minimized, while it also reduces background and facilitates multiplexing by barcoding. Here, we implement barcodes for up to 448 genes per color channel and demonstrate two-color imaging. Furthermore, electrophoresis increased the transfer efficiency and reduced lateral diffusion, when compared to RNA transfer by passive diffusion (as used by e.g. Visium^9^), resulting in a more faithful RNA blot that retains cellular resolution.

We applied EEL to a full sagittal section of the mouse brain where 440 genes were measured in little over two days of imaging, which enabled the study of spatial regions, gradients and borders defined by gene expression. Moreover, the cells could be segmented to yield the spatial transcriptome profiles of more than 128,000 single cells, enabling clustering and visualization of cell type-specific expression profiles in their spatial context. To further demonstrate the robustness, scalability and versatility of EEL, we then created a transcriptome atlas of the mouse brain comprising seven additional sagittal sections, where we examined spatial organization. Finally, we applied EEL to the human visual cortex and found that EEL greatly reduced the highly autofluorescent lipofuscin deposits that normally restrict RNA detection by smFISH in human tissue.

To aid implementation of EEL in other labs, we provide a detailed description of the hardware — schematics, parts list, build instructions — together with all the necessary software for instrument control, image processing, data analysis and visualization.

## Results

### EEL protocol

We developed and optimised a protocol to transfer RNA from a tissue section onto a capture slide with high efficiency and minimal spatial distortion, by actively forcing the RNA onto the surface through electrophoresis. Briefly (see Methods for a detailed protocol), the RNA capture slide is a glass coverslip coated with an optically transparent and electrically conductive layer of indium tin oxide (ITO), which is modified with oligo-dT and positively charged poly-D-lysine to capture RNA both by hybridization and electrostatic attraction (Fig. 1a). A 10 μm cryosection is placed onto the capture slide and stained nuclei are imaged for cell segmentation. The tissue is permeabilized and an electric potential difference of 10 V/cm is applied for 20 minutes, where the capture slide acts as the anode (Extended Data Fig. 1c). When the RNA transfer has been completed, the tissue sample is digested, leaving only the captured RNA on the surface. Tissue removal speeds up the subsequent detection chemistry because reagents and probes do not need to diffuse through the tissue.

**Fig. 1.**
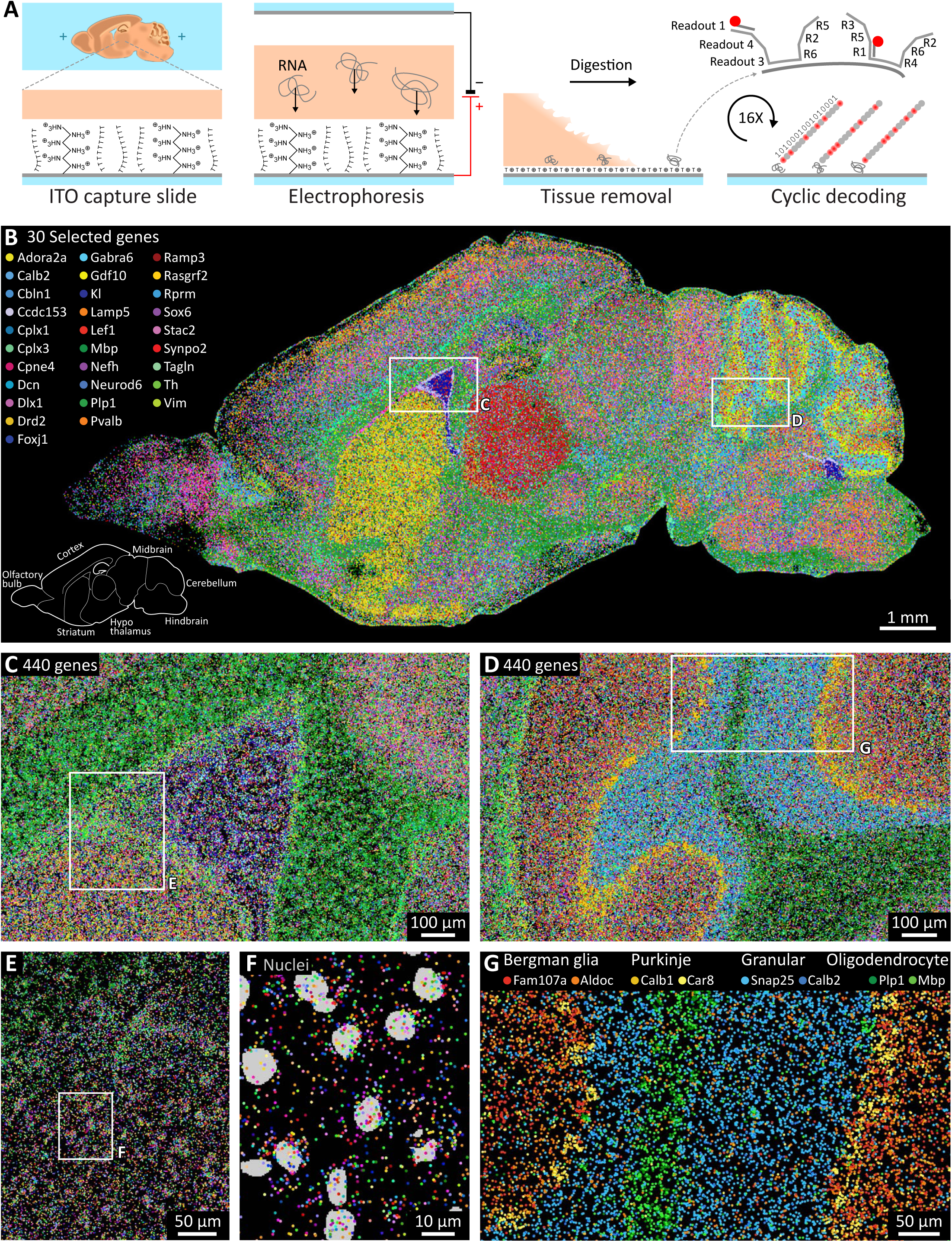
EEL method and 440 gene mouse dataset. **a**, Schematic illustration of the EEL protocol, including RNA transfer by electrophoresis, capture on the ITO slide, tissue removal and cyclic fluorescent decoding. **b**, Results of EEL on a sagittal mouse brain section, where every dot is a single molecule of RNA belonging to one of the 30 selected genes to highlight large anatomical structures. The total experiment contained 440 genes. **b, c**, Zoomed in results for all genes around lateral ventricle (**c**) and cerebellum (**d**). High resolution imaging (<500 nm) yields sub-cellular resolution (**e, f**) and cell type marker expression patterns for small anatomical features (**g**).

Next, to detect up to 448 species of RNA per color channel, we designed a set of binary codes with 6 positive bits out of 16 bits total, and minimum 4 bits of difference between any pair of codes. For each desired target gene, a set of highly specific encoding probes was designed to tile the transcript, each carrying overhanging tails encoding the 6 positive bits of the selected barcode (Fig. 1a).

For each experiment, the full set of encoding probes for all target genes was pooled and hybridized upfront and was then detected by 16 cycles of fluorescent readout probe hybridization to the tails, imaging, and TCEP-mediated fluorophore cleavage. The entire process was implemented on a custom-built open-source fluidic system, integrated with a commercial microscope to perform the barcode detection automatically (Fig. 1a, Extended Data Fig. 1d-f).

As proof of principle, we targeted 440 genes and 8 empty-barcode controls, which after image processing and barcode decoding, resulted in 8,871,209 detected RNA molecules with a false positive rate of 0.025% ± 0.011% per gene. The spatial expression patterns could be analysed both at the scale of the full tissue section (Fig. 1b), of smaller anatomical structures (Fig. 1c,d,g) and at the single-cell level (Fig. 1e,f).

### EEL transfer efficiency and sensitivity

We estimated the RNA transfer efficiency by comparing RNA numbers detected by smFISH directly after transfer and tissue digestion, with those of a consecutive tissue section processed using regular in-tissue smFISH. We found that around 19% of the RNA was transferred onto the surface (Fig. 2a, Extended Data Fig. 2). However, barcode detection and decoding caused an additional loss of detected transcripts. Comparing the non-barcoded osmFISH dataset in mouse somatosensory cortex^2^ with EEL for the same genes showed that the complete end-to-end EEL protocol achieved a 4.4% detection efficiency. One key to RNA capture efficiency was the addition of poly-D-lysine to the surface, which enhanced the transfer efficiency 60-fold compared to capture by oligo-dT alone (Extended Data Fig. 2a,b).

**Fig. 2.**
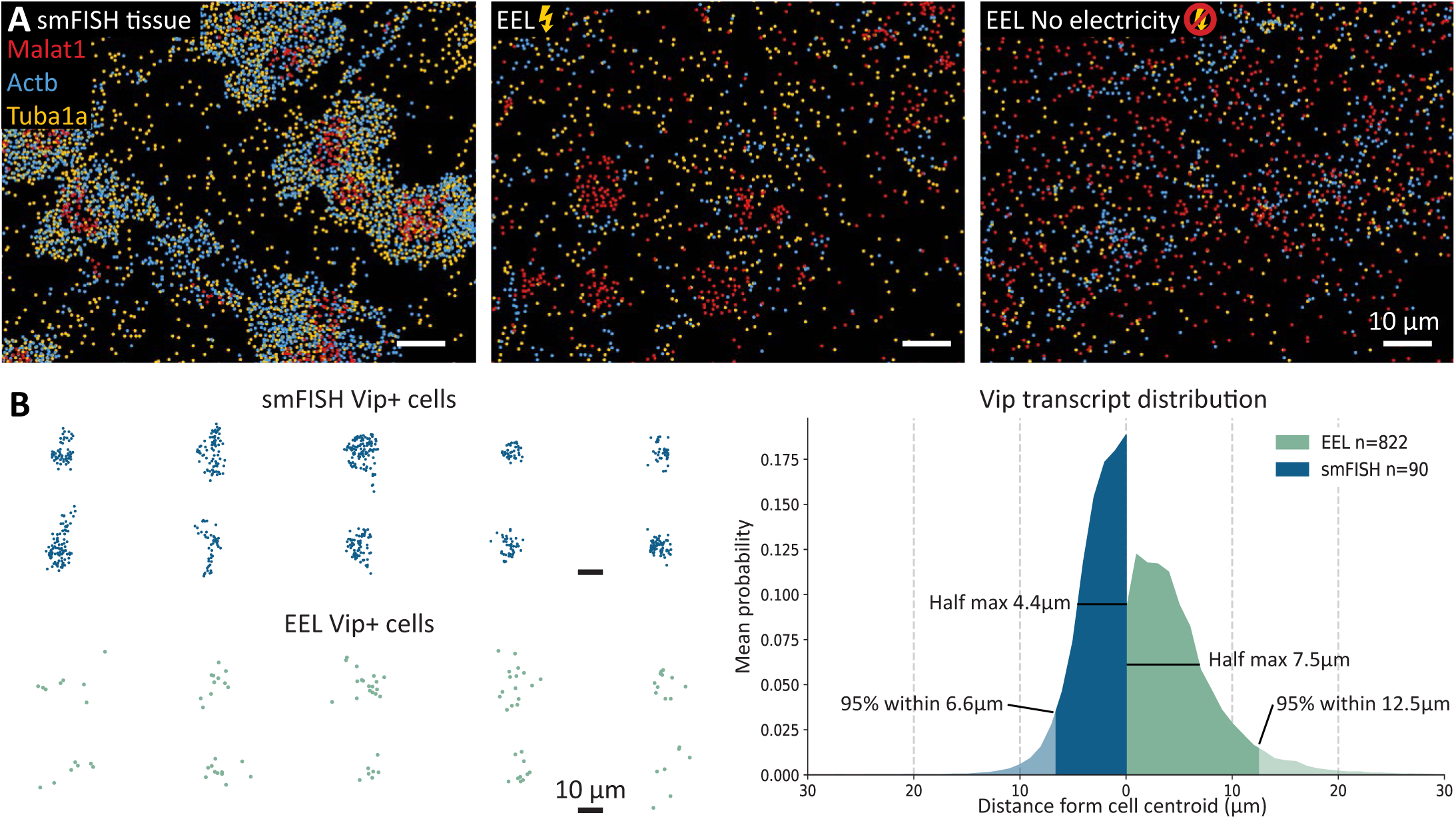
Electrophoresis enhances RNA blot accuracy. **a**, 3 gene smFISH in tissue (left), EEL with electrophoresis (middle) and EEL without electrophoresis (right). With electricity the EEL RNA distribution looks more similar to the original tissue situation than without it. **b**, Distribution of Vip RNA in tissue smFISH and EEL, indicating that samples processed with EEL show little lateral diffusion from cell centroid.

Because the capture surface was also the electrode, we were concerned that electrolysis of water at the anode would lead to a local drop in pH that would interfere with the capture by neutralizing or reversing the charge of the RNA molecule. To verify that pH did not drop, we directly measured surface pH by sandwiching particles doped with the pH-sensitive fluorophore pHRodo between the slide and tissue section, which demonstrated that the pH was maintained during electrophoresis (Extended Data Fig. 3).

### Lateral diffusion

Electricity reduced lateral diffusion during RNA transfer both on a large and single-cell scale, so that the original tissue organization was preserved by the tissue blot (Extended Data Fig. 4a,b). The distinct cellular pattern of RNA reflecting the sparse localization of cell bodies in the mouse brain was also better preserved when electricity was used (Fig. 2a, Extended Data Fig. 2d, 4c, 8). To quantify the preservation of spatial structure, we used Ripley’s L function^23^ which measures deviation from homogeneity. We found a stronger peak for the typical cell size (∼10 μm) for conditions that included electricity — as compared with those that did not — indicating that electrophoresis better preserved the original spatial distribution (Extended Data Fig. 2c).

As an independent and more direct estimate of the extent of lateral diffusion, we exploited the sparse distribution of cortical interneurons that express neuropeptides at high levels. We identified individual cells expressing *Vip* and measured the distance from the cell centroid to nearby *Vip* transcripts. For EEL data, the full width at half maximum was 7.5 μm, compared to 4.4 μm in tissue using smFISH (Fig. 2b). Thus, the lateral diffusion under EEL conditions was smaller than one cell diameter on average.

### Barcoding and image acquisition

Capturing RNA on a surface speeds up imaging by reducing or eliminating the Z-stack, and improves the signal-to-noise ratio by diminishing background noise normally caused by tissue autofluorescence and scattering. Together with improvements to the microscope setup, these factors increased the imaging throughput 40-fold compared to osmFISH^2^, reducing the image acquisition time to 76 seconds per mm^2^, compared to 51 minutes/ mm^2^ for osmFISH. Additionally, EEL encodes up to 448 genes per channel in 16 rounds, compared to a single gene per channel and round in osmFISH, resulting in another 28-fold increase in throughput.

Altogether, these improvements meant that a complete 448-gene EEL experiment covering 1 cm^2^ of tissue could be completed in 58 hours, many orders of magnitude faster than osmFISH. The EEL protocol was also nearly fully automated, leaving only four hours of hands-on processing time.

All datasets reported here were collected with one color channel, but the number of genes can simply be scaled by adding more channels, which is time-efficient because it only requires an additional exposure but no other additional steps. As proof of principle, we demonstrated this by measuring 883 genes in a human glioblastoma sample using two sets of encoding probes corresponding to detection probes carrying Alexa-647 and Cy3 dyes, which resulted in the detection of 883 genes over 87 mm^2^ in 71 hours (Extended Data Fig. 1g).

### Data analysis

smFISH-based spatial transcriptomics methods generate image datasets with the size of several terabytes. To efficiently process this data, we developed pysmFISH, an open-source distributed computing pipeline that automatically performs background filtering, RNA detection, alignment and decoding of barcoded, as well as sequential smFISH experiments (Extended data Fig. 5).

Furthermore, to facilitate the interactive exploration of the resulting point clouds that can contain many millions of molecules per tissue section, we developed FISHscale, which is a Python-based 3D visualisation and analysis tool for single or multiple large-scale point-based spatial transcriptomic datasets.

FISHscale leverages GPU acceleration to enable dynamic zooming, panning and 3D rotation at high frame rates even with tens of millions of dots.

### Regions and borders

Highly multiplexed spatial transcriptome datasets enable the automatic generation of atlases of complex tissues^2,24,25^. We binned RNA spots in a hexagonal grid and performed principal component analysis that could be clustered to generate a regionalized map of the tissue that recapitulated known anatomical structures in the mouse brain (Fig. 3a, Extended Data Fig. 8a-d).

**Fig. 3.**
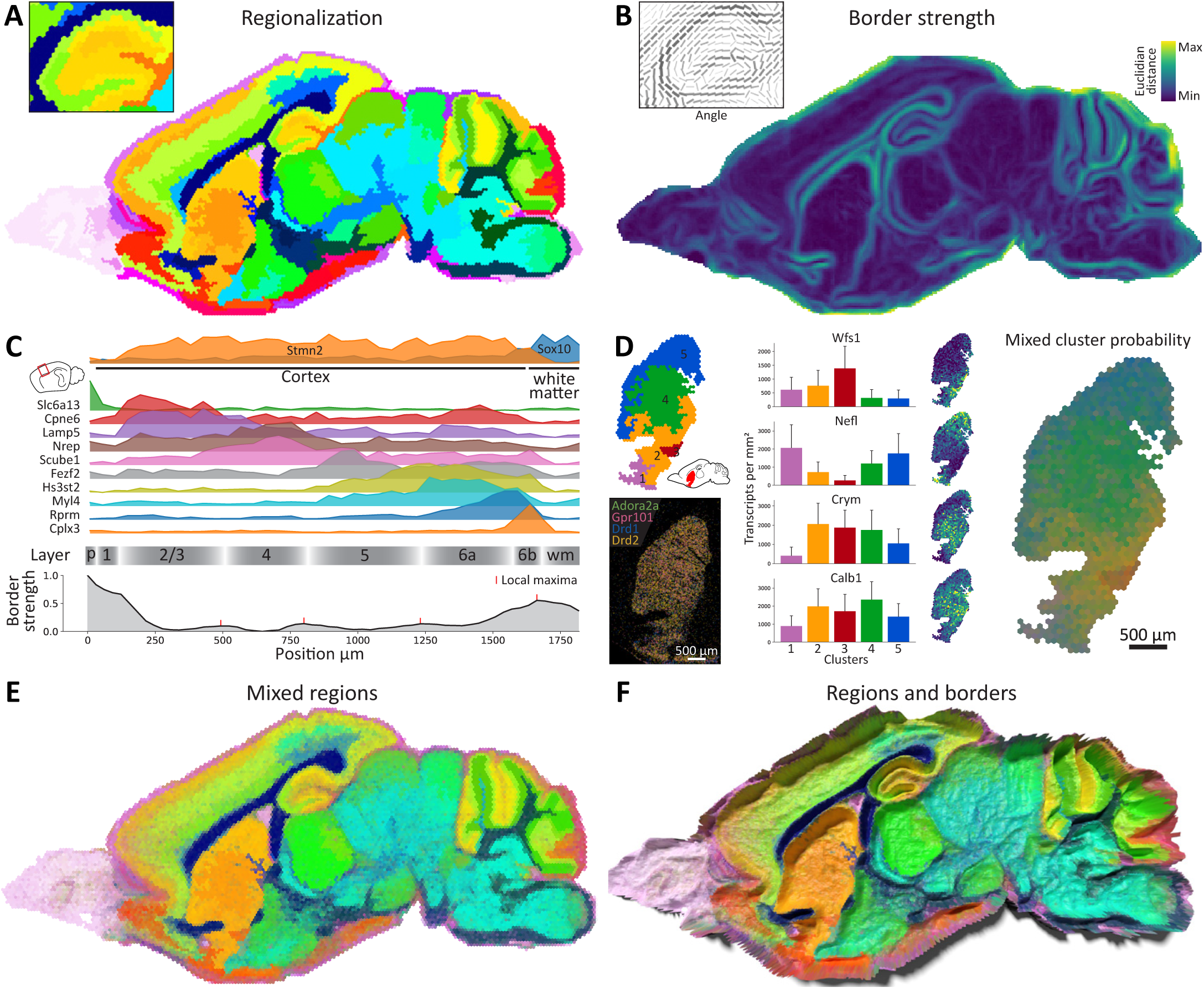
Data-driven anatomical regions and borders. **a**, Clustering of hexagonally binned expression data results in distinct anatomical regions. **b**, Local transcriptional heterogeneity can be quantified to obtain border strength and angle. **c**, Transcript densities for selected genes show layer-restricted expression between pia and white matter, where interfaces between domains are marked by local maxima in border strength. **d**, The striatum has no clear sub-regions but contains spatial gradients of certain genes. Mixing region label colors by cluster probability of cluster identity displays these gradients better than discrete colors. **e**, Mixed cluster colors for the full sample show gradients as well as discreet borders. **f**, Mixed region colors rendered with border strength as height to show difference between sharp borders and gradients.

To complement the regionalization, we introduced a border sharpness metric by calculating the local directional dissimilarity as a measure of the strength and orientation of the potential borders (Fig. 3b, Extended Data Fig. 6e). We observed the strongest transcriptional difference between grey and white matter, between the cell-dense and cell-sparse regions of the hippocampus and cerebellum and surrounding the striatum and thalamus (Fig 3b). Weaker, but clear borders were observed between the layers of the cortex, where genes showed varying layer-specificity (Fig. 3e, Extended Data Fig. 6f).

In contrast, the striatum contained no clear sub-regions but instead showed multiple gradients^26^ resulting in arbitrary splits with a clustering approach (Fig. 3d) (See also Partel *et al*. (2020)^25^). To model such gradients, we trained a classifier on the cluster labels, and then mixed the original region colors based on the class probability of each cluster label (Fig. 3d). This way of visualisation preserved highly distinct regions that showed sharp borders, as well as gradients, and could be jointly visualised with border strength (Fig. 3e,f).

### Single-cell resolution

Assigning expressed RNA molecules to cells enables analysis of cell types and cell states in their spatial context, but requires segmentation of cell bodies^7,27,28^. To generate single-cell expression profiles from EEL data, images of propidium iodide-stained nuclei taken at low magnification before tissue digestion were segmented with Cellpose^29^, expanded and registered to the RNA signal with the help of fiducial markers so that molecules could be assigned to cells (Fig. 4a, Extended Data Fig. 7, Methods). After quality control, this resulted in an expression matrix of 128,813 single cells and 440 genes, with a total of 6,470,942 transcripts assigned to cells (73% of all molecules detected in this tissue section).

**Fig. 4.**
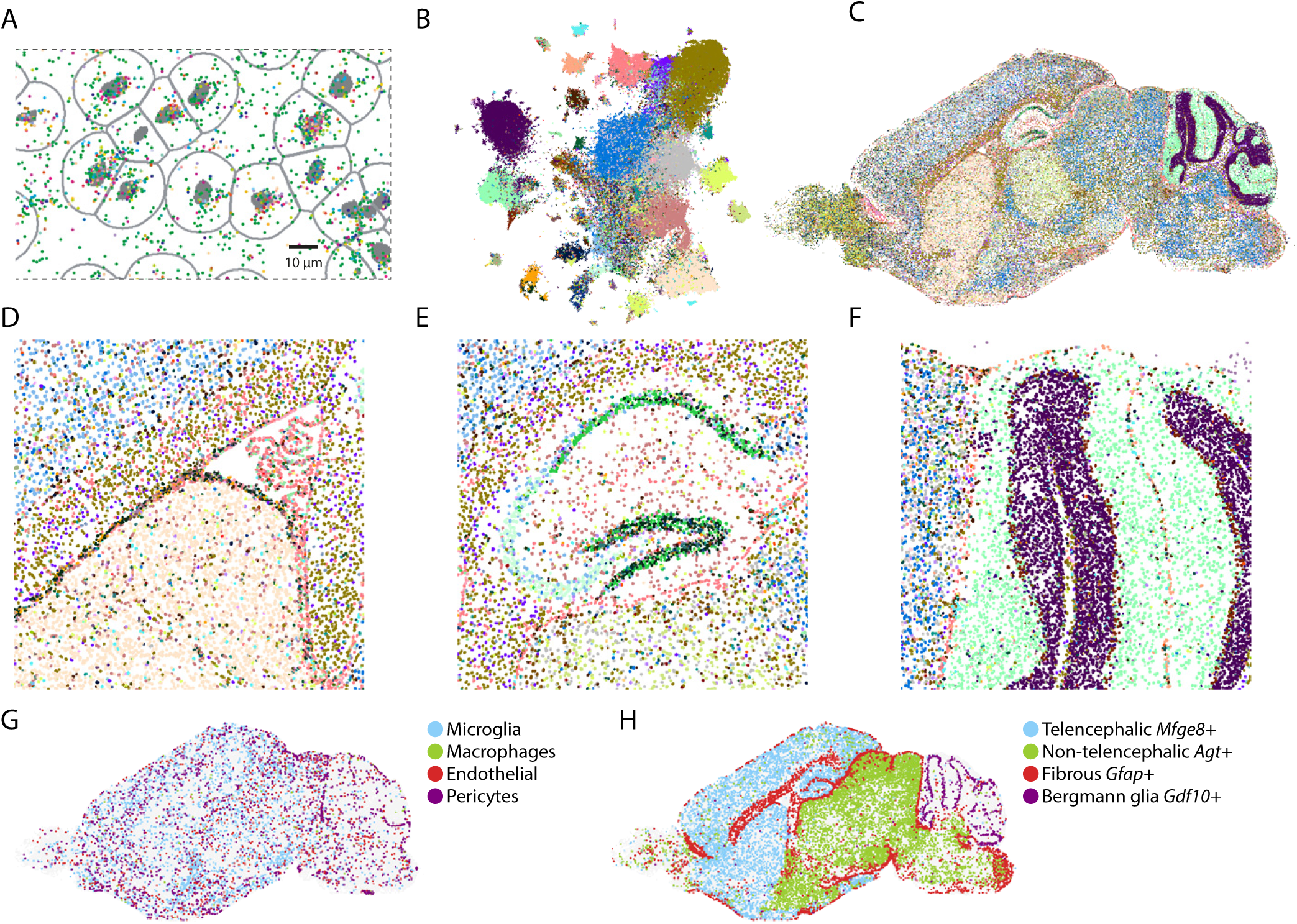
Single cell analysis of EEL data. **a**, RNA spots assigned to cells inside the expanded segmentation masks of aligned nuclei (grey borders). **b**, t-SNE of single-cell profiles where colors indicate the 147 clusters. **c**, Spatial cell type map where every dot is a single cell colored by cluster identity as in (b). **d-f**, Magnified views of (c) showing the ventricle including rostral migratory stream (d), the hippocampus (e) and the cerebellum (f). **g**, Same as (c) but showing only microglia, macrophages, endothelial cells and pericytes. **h**, Astrocyte types showing distinct spatial distributions.

Next, we applied a standard single-cell clustering pipeline (see Methods) to construct a spatial map of cell types in the mouse brain, resulting in 147 distinct clusters representing major cell types (Fig. 4b-f). Examining these clusters, we found both abundant cell types with little spatial structure such as microglia and endothelial cells, as well as highly spatially structured neurons and glia (Fig. 4g,h). For example, several distinct types of astrocytes occupied largely non-overlapping domains (Fig. 4h): olfactory bulb, the rest of the telencephalon, cerebellum (Bergmann glia) and the non-telencephalic brain. The *Gfap+* fibrous astrocyte subtype was found in white matter and covered most of the brain surface (glia limitans) with the exception of the cerebellum. Cortical layer-specific excitatory and inhibitory neurons were observed, as well as distinct hippocampal neurons of the CA1, CA3, dentate gyrus and the molecular layer interneurons (Fig. 4g). Several clusters corresponded to migrating neuroblasts of the rostral migratory stream into the olfactory bulb and of the dentate gyrus subgranular zone (Fig. 4d-e, Supplementary Fig. 8).

These results demonstrate the power of EEL to reveal the cell type composition of the mouse brain from a single experiment.

### Mouse brain atlas

Next, to demonstrate the robustness and scalability of EEL, we generated an atlas of the mouse brain by measuring the expression patterns of 168 genes in seven sagittal sections starting at the midline and moving laterally with a spacing of ∼600 μm, detecting a total of 17,151,357 molecules (Extended Data Fig. 9). We observed region-specific gene expression of various structures, which all showed clear correspondence between consecutive sections (Fig. 5a). This enabled the regionalization of individual sections and linking the resulting regions in 3D (Extended Data Fig. 10). Interestingly, our data showed in great detail the sharp border between telencephalic and non-telencephalic astrocytes, marked by *Mfge8* and *Agt*, respectively^1^. Having access to multiple sagittal sections confirmed that the border closely followed the telencephalon-diencephalon divide, further reinforcing the notion that these astrocyte types are developmentally specified.

**Fig. 5.**
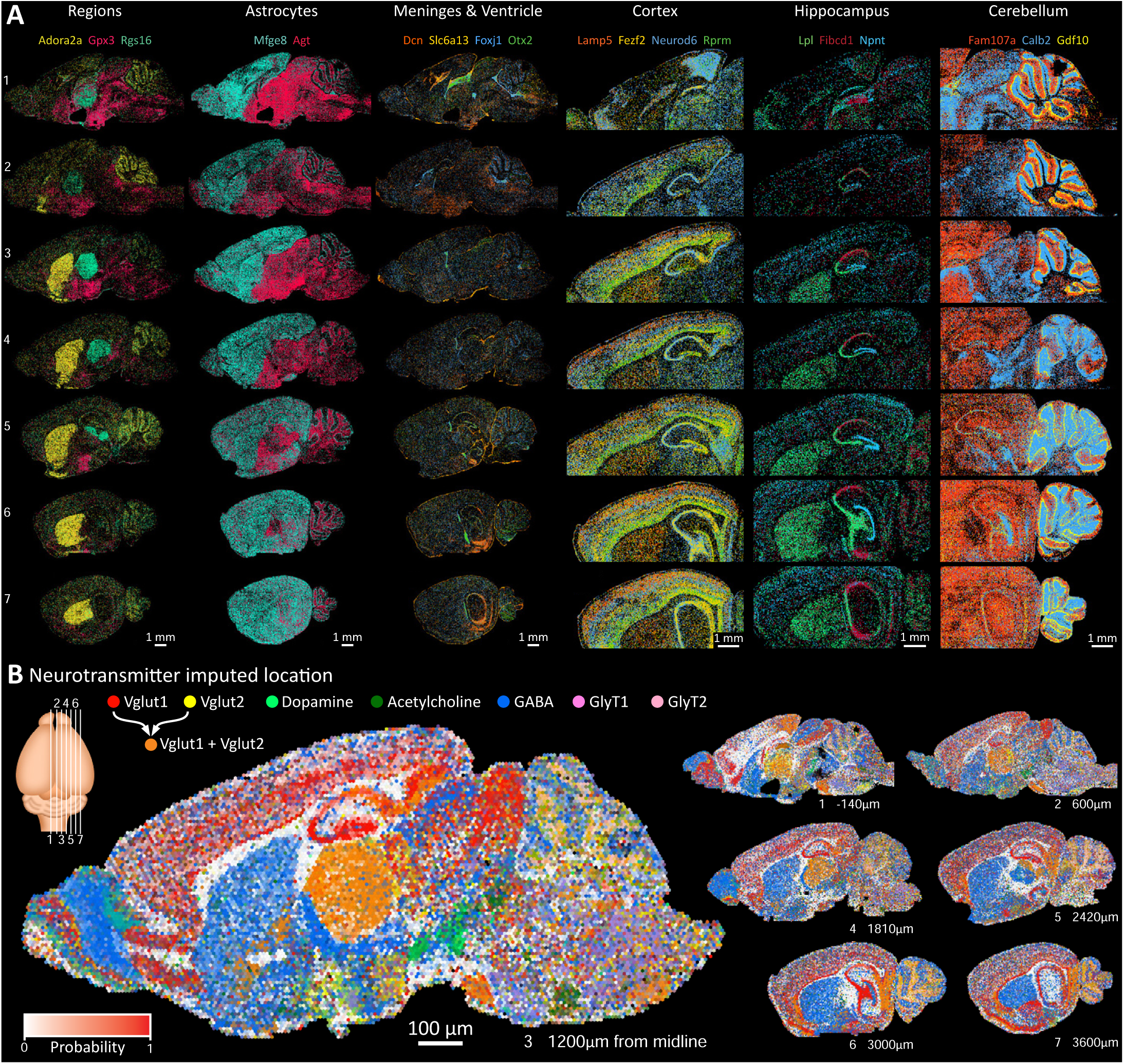
Mouse brain atlas of 7 sagittal sections. **a**, Raw RNA signals for selected markers to highlight various anatomical structures. From left to right: Markers for striatum (Adora2a), thalamus (Rgs16) and hypothalamus (Gpx3); Mfge8 labels telencephalic astrocytes while Agt labels non-telencephalic astrocytes; markers for meninges (Dcn, Slc6a13), ependymal (Foxj1) and choroid plexus epithelium (Otx2); layers of the cortex; Hippocampal CA1 (Fibcd1), CA3 (Lpl) and dentate gyrus (Npnt); cerebellar Bergman glia cell body (Gdf10), processes (Fam107a) and granule cells (Calb2). **b**, Probabilities of neurotransmitter location by imputing single-cell RNA-seq data with the spatial mouse atlas, shows neurotransmitter domains. Intensity reflects probability of neurotransmitter identity and colors are proportionally mixed where neurotransmitters overlap.

High multiplexing not only facilitates the study of many regions and cell types in a single experiment, but can also be used to spatially embed single-cell RNA-seq data. We used a generalised version of Tangram^30^, named Bonefight^31^, to align our previously published single-cell census of the mouse brain^1^ with the spatial mouse atlas, resulting in a putative anatomical location for all cell types, largely in agreement with expected locations (Extended Data Fig. 11). Once cell types have been aligned, their properties can be transferred to the spatial domain. For example, we generated spatial maps of neurotransmitter usage by summing the spatial probabilities of cell types that share a specific neurotransmitter and projecting them spatially (Fig. 5b, Extended Data Fig. 12). Overlaying all neurotransmitters in the same section showed the regional preference of one, or sometimes multiple neurotransmitters. For example, *Vglut1* and *Vglut2* were both present in the thalamus, subiculum, and the pontine grey (Fig. 5b, orange).

### Human brain atlas

The study of human brain samples by spatial methods has been limited both by the size of the human brain and by the presence of lipofuscin, an age-related accumulation of highly auto-fluorescent lipid-containing lysosomal residue which is mostly found in neurons. The presence of lipofuscin has precluded us from studying the human brain in the past using osmFISH because the strong auto-fluorescence interfered with the weak fluorescence signal from individual mRNA molecules. In contrast, we found that the EEL protocol eliminated most of the lipofuscin and thereby reduced autofluorescence, thus enabling the study of human samples (Extended Data Fig. 13).

We applied EEL on a 0.75 cm^2^ section of the human primary visual cortex and measured the expression of 445 genes (Fig. 6). Individual genes allowed us to identify region-specific expression patterns repeated along the entire cortical structure: glial and other non-neuronal cell types (Fig. 6i), cortical superficial and deep layer-specific expression (Fig. 6ii, 6iii), and spatial distribution of inhibitory cell types (Fig. 6iv, 6v).

**Fig. 6.**
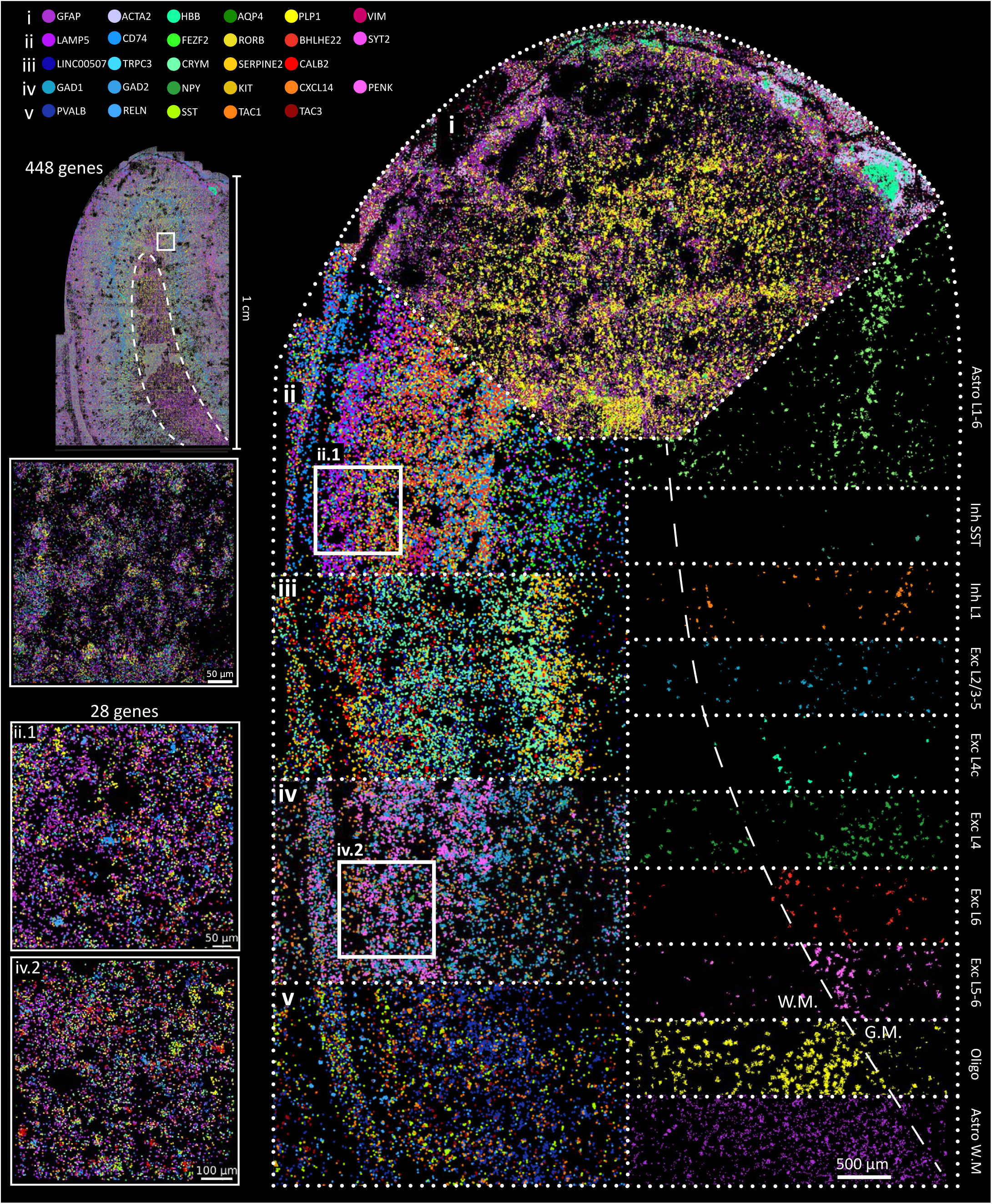
EEL results on adult human visual cortex. Highlighting the RNA spots of 28 genes in the primary visual cortex to show: (i) markers for non-neuronal cell types; (ii) superficial layer neuronal markers; (iii) deep layer neuronal markers; (iv, v) and inhibitory neuron markers. On the right side, GraphSAGE molecule embedding clusters show cell types spatially organized in anatomical compartments, such as layers (L), white matter (W.M.) and gray matter (G.M.).

To explore the spatial structure of the human visual cortex, we trained an unsupervised graph neural network (GNN) constructed on RNA molecule neighbourhoods (as proposed by Partel & Wälby, 2021)^32^. We found that molecule embeddings corresponded to spatially and molecularly distinct domains, and we observed that some of these domains corresponded to known cell types of the human cortex, such as excitatory and inhibitory neurons, astrocytes, and mature oligodendrocytes (Fig. 6, right). The human visual cortex differs from other cortical areas by the characteristic band of Gennari^33^, which contains axonal tracts carrying visual input from the thalamus, that we could identify along layer 4 by both the spatial location and molecular profile in our data (Fig. 6 L4c).

These findings demonstrate that EEL can be successfully applied to human adult brain tissue, as the high level of multiplexing and spatial resolution allowed us to characterise the distinct molecular profiles of cells in the cortex that correspond to cell types or states (Extended data Fig. 14).

## Discussion

Here, we have described a spatial transcriptome profiling method that is capable of high multiplexing, high spatial throughput and high resolution, thus addressing the difficulties in elucidating the spatial distributions of large numbers of cell types within complex tissues and enabling the study of entire sections of mouse brain in just two days of imaging. Additionally, by reducing the detrimental effect of lipofuscin to imaging, EEL enables the investigation of human samples at meaningful scale.

EEL builds on concepts from both microscopy- and sequencing-based spatial transcriptomic methods. Similar to sequencing-based methods^9–15^, we transferred the RNA to a surface, although EEL introduced an active RNA transfer step instead of relying on passive diffusion. We found that electrophoresis better preserved the spatial tissue organisation on cellular scale. Similar to microscopy-based methods, we then detected mRNA molecules *in situ* using targeted probes^34^.

The main advantage of sequencing-based methods is that they can capture and sequence the full transcriptome. In contrast, EEL — like most micros-copy-based techniques — relies on panels of selected probes, albeit with a flexible design that potentially is scalable to thousands of genes. For example, with five color channels EEL could detect more than two thousand genes, similar to the typical number of variable genes found in single-cell RNAs-seq experiments. With additional imaging cycles and modified barcode schemes, EEL could be scaled further, as has been previously demonstrated for surface-based RNA detection with smFISH (e.g. 10,212 genes, RNA SPOTs^34^).

Furthermore, EEL is an order of magnitude less expensive than sequencing-based methods at $600 per experiment (∼$0.005/cell). Additionally, EEL has the advantage that the capture surface is continuous so that there are no gaps between capture features. The diffraction-limited imaging resolution of 200-400nm enables single-cell transcript assignment through cell segmentation, resulting in a true cell-by-gene data matrix.

As trade-off for high spatial throughput, EEL showed a lower sensitivity compared to tissue-based smFISH methods, currently considered the most sensitive methods for RNA quantification. Nevertheless, the data quality was similar to typical single-cell RNA-seq data and allowed for the spatial analysis of both transcriptionally defined regions and their borders, as well as constructing atlases of cell types at single cell level. With future improvements to RNA stability and barcode detection, EEL’s sensitivity can be further improved.

EEL makes it possible to analyze whole mouse organs, as shown here for the mouse brain. However, even with a greatly increased imaging speed, further improvements would be needed to be able to image whole human organs. One factor limiting imaging speed in our current setup was a slight curvature of the capture slide caused by the flow cell design, that we compensated for by imaging a small Z-stack. By designing a more rigid flat flow cell and further refinements to our fluidics timing, it should be possible to scale EEL beyond ten cm^2^, approaching the size of many important human brain structures such as the midbrain, hindbrain, and cortical lobes.

Early gene expression atlases (e.g. the Allen brain atlas^35^) have served as immensely valuable resources for the wider research community. However, enormous resources and complex organisation were required to generate such datasets, with the result that they could not be applied to many subjects, diseased tissues or genetic animal models. With the development of highly multiplexed spatial transcriptomics — scalable to large tissue areas and with rapid and robust automation — it now becomes possible to generate bespoke atlases for specific research questions. EEL enables the study of complex tissue specimens at scale and facilitates the study of the molecular organisation of the human brain.

### Online content and data availability

Data: http://mousebrain.org/

Supplemental table: Excel file with all probes.

Protocol:

https://www.protocols.io/view/eel-fish-t92er8e

ROBOFISH:

https://www.protocols.io/view/robofish-construction-bcrciv2w,

https://github.com/linnarsson-lab/ROBOFISH

FISHScale:

https://github.com/linnarsson-lab/FISHscale

pysmFISH:

https://github.com/linnarsson-lab/pysmFISH_auto

Olygopy:

https://github.com/linnarsson-lab/oligopy

## Acknowledgements

We would like to thank Alex K. Shalek, Andrew W. Navia, David Poxson, David Nilsson and Daniel Simon for valuable advice on ITO cleaning and functionalization. We thank Long Cai and Chee-Huat L. Eng for sharing probe samples to set up barcoding. We would like to thank Dirk Keene for generously providing human brain specimens. We thank Amit Zeisel, Gioele La Manno and all members of the Linnarsson lab for fruitful discussion of the project. We thank P. Lönneberg for maintenance of the computing cluster and A. Johnsson for project management. This work was supported by grants from the Swedish Foundation for Strategic Research (SB16-0065), the Knut and Alice Wallenberg Foundation (2015.0041, 2018.0172, 2018.0220), US National Institutes of Health (NIH) grant U01 MH114812-02, the Erling-Persson Foundation (HDCA) and the Torsten Söderberg Foundation to S.L.

## Author contributions

L.E.B., S.C. and S.L. conceived the study. L.E.B., A.M.A., C.M., J.J., C.Ö. and D.F.G. optimized the protocol. L.E.B performed mouse experiments. A.M.A. performed human experiments. S.C. developed and performed image analysis. L.E.B, A.M.A. and S.L. performed data analysis. L.E.B. build the ROBOFISH system. A.M.A. and S.L. designed the probes and barcoding. R.H. and E.L. provided human samples. L.E.B., A.M.A., S.C., and S.L. wrote the manuscript. All authors provided critical feedback and helped shape the research, analysis and manuscript.

## Competing interests

The authors declare the following competing interests: L.E.B., A.M.A., C.M., J.J., D.F.G., S.C. and S.L. are shareholders in EEL Transcriptomics AB, which owns intellectual property rights to the EEL method, including a patent application with L.E.B., S.C. and S.L. as co-inventors. S.L. is a paid scientific advisor to, and S.C. is a current employee of Rebus Biosystems, which has licensed the EEL intellectual property rights. The remaining authors declare no competing interests.

## Methods

### Protocol

The full step-by-step protocol is available online at https://www.protocols.io/view/eel-fish-t92er8e.

### Sample collection

Animal handling and tissue harvesting methods followed the guidelines and recommendations of local animal protection legislation, and were approved by the local committee for ethical experiments on laboratory animals (Stockholms Norra Djurförsöksetiska nämnd, Sweden, N 68/14). Two wild-type CD1 female mice postnatal day 41 were used for the mouse atlas and mouse 448 gene experiments. Mice were transcardially perfused with ice cold oxygenated artificial cerebrospinal fluid solution. Brains were harvested, submerged in Tissue-Tek optimum cutting temperature (O.C.T. Sakura) and snap frozen in a slush of iso-pentane (Sigma) and dry ice, before storage at-80°C.

Human tissue collection was performed after obtaining permission from the decedent’s next-of-kin as previously described by Hodge *et al*. (2019)^36^, in accordance with the provisions of the United States Uniform Anatomical Gift Act of 2006 described in the California Health and Safety Code section 7150 (effective 1/1/2008) and other applicable state and federal laws and regulations, and with ethical approval from the Swedish Ethical Review Authority (2019-03054 for human adult brain, 2020-02096 for human glioblastoma). Human serological screening for infectious disease (HIV, hepatitis B and hepatitis C) was conducted using donor blood samples, and only considered if it was negative for all three tests. Human cortex V1C was sliced, submerged in Tissue-Tek optimum cutting temperature (O.C.T. Sakura) and snap frozen in a slush of iso-pentane (Sigma) and dry ice before storing it at-80 °C.

### Capture slide preparation

ITO coverslips (24 × 60 mm #1.5 thickness, Diamond Coatings Ltd.) with a surface resistivity of 30-60 Ω/ square were cleaned by three successive washes of 20 minutes in a beaker glass filled with acetone (Sigma), iso-propanol (Sigma) and dH_2_O (Thermo) placed in a Ney ULTRAsonik 28X sonicator set to maximum power. Slides were stored in dH_2_O and used between 5 and 30 days after production. To functionalize the surface, the slides were placed in an Epridia E103 rack (Fischer Scientific), dried with nitrogen gas and submerged in a 2% (v/v) solution of (3-Glycidyloxypropyl)Trimethoxysilane in acetone for 2 hours under a nitrogen atmosphere. Coverslips were rinsed once with acetone and dried with nitrogen gas. Then the oligo-dT mixture consisting of 10 μM 5’ amine-modified oligo-dT-60 (/5AmMC6/ UUUUGACTCGTTTTTTTTTTTTTTTTTTTTTTTTTTTTTTT/ iSuper-dT/TTTTTTTTTTTTTTTTTTTTTTTTTTT/iSuper-dT/ TT, IDT) in 1X Schott Spotting Solution was prepared. The ITO coated side of the coveslip was identified using a multimeter, 40 μl of the oligo-dT mixture was placed on the centre and covered with a 24 × 24 mm plastic Hybrisilp (Grace Biolabs). The oligo-dT were let to react with the epoxy groups for 1 hour at 25 °C. Afterwards, the coated slide was washed 5 times with SSC 2X (Sigma) followed by 2 washes with dH_2_O. Remaining epoxy groups were blocked for 30 minutes at room temperature with a 0.1% (w/v) solution of poly-D-Lysine (MW 70,000 - 150,000 (Sigma)) in dH_2_O, followed by 3 washes with dH_2_O. Europium-doped beads with a diameter of 0.2 and 1 μm (Thermo, 1 μm custom produced) were each diluted 1:333 in dH_2_O and deposited on the surface by placing a 100μl drop on the coated area for 3 minutes. The bead mixture was pipetted off and slides were dried to air.

Coated capture slides can be stored for at least 2 days under nitrogen atmosphere. Cryosections of 10 μm were cut and captured on the coated area, then dried for a few minutes and stored at –80 °C until use. Frozen slides with tissue sections can be stored for a few months before proceeding to the transfer step.

### Probes

Direct labelled smFISH probes for *Malat1, Actb* and *Tuba1a* were obtained from Biosearch Technologies. Detection probes for barcoded EEL experiments were obtained from IDT and consist of a 5’-conjugated Alexa Fluor 647 dye, followed by a thiol linker (/iThioMC6-D/) and then the 20 nucleotide long detection sequence (Supplementary table).

The RNA-binding sequence of the encoding probe design was based on previously reported methods^18,37^ and implemented in the software package Oligopy (For a similar tool see PaintSHOP by Hershberg *et al*. (2021)^38^). Each probe consisted of a 26-32 nucleotide long sequence, reverse complementary of the target RNA sequence, and 1 or 2 overhanging tails containing the 6 detection sequences of 20 nucleotides each, to which the detection probes can bind. These 6 sequences were added to the RNA binding sequence in random order to reduce the potential steric hindrance effect on signal intensity when they accumulate over sequential rounds of hybridization. Furthermore, readout tails were separated by two nucleotides (alternating TT, AT, TA and AA).

RNA binding regions were selected to have a Gibbs free energy (ΔG) at 37 °C of -28 kcal/mol or less, GC content of 40-60% and a maximum ΔG at 37°C for hairpin or homodimer formation of -9.0 kcal/mol. Then, probe specificity was analysed using BLAST, and probes with more than 60% identity to five or more off-target RNA species were dropped. Finally, sets of maximum 28 probes that would tile with a minimum gap of 2bp on the RNA, were retrieved, and genes that did not reach a minimum of 10 probes were dropped.

To prevent optical crowding, we optimised the distribution of genes over the available barcodes using prior knowledge from single-cell RNA seq data. Whenever possible, the genes expressed in the same cell type were not labelled in the same decoding cycle.

Encoding probes were ordered either from IDT as oPools (mouse atlas experiment) at ready-to-use concentrations, or as Twist Bioscience Oligo Pools for the 448 gene mouse and human experiments.

For the IDT oPool probe sets, the fraction of full-length oligonucleotides was expected to be only ∼40% for the 186 nt long probes, because of truncation during chemical DNA synthesis. Placing 2 tails at both 5’ and 3’ of the RNA-binding sequence (as introduced by MERFISH^18^, would potentially result in truncated probes retaining RNA binding ability but lacking some of the redout sequences. To avoid the issue, the entire RNA-binding sequence was placed on the 5’ side of the oligo as one tail, so that only full length probes could bind the RNA.

For higher-complexity probe sets, Twist Bioscience Oligo Pools were used and amplified using a previously published protocol^39^ with the following adaptations. We performed an initial PCR amplification to generate an intermediate stock and used it as the template for subsequent experiment, not to exhaust the original probe pool. Then, the probes were amplified again by PCR, followed by *in vitro* transcription, after which the RNA was purified using the Zymo Oligo Clean and Concentrator. Reverse transcription was performed using a 5’-modified forward primer with an amine group so that the encoding probe could be fixed by paraformaldehyde (PFA) for better signal stability. Lastly, RNA was degraded by alkaline hydrolysis and ssDNA probes were purified. Probe sequences are available as Supplementary table.

### EEL

Washes were generally performed by pipetting 200 μl of solution onto the sample or, in the cases where a flow cell was used, the internal volume of the flow cell was at least replaced twice for one wash.

Slides with tissue were thawed and two reference crosses were drawn on the bottom of the coverslip flanking the tissue sample with a solvent-resistant Moist Mark Plus pen (Cancer Diagnostics Inc., Fig. 1a). Nuclei were labelled for 5 minutes with a 1 μg/ ml solution of Propidium Iodide (Sigma) in 2X Sodium Sodium-Citrate buffer (SSC, Sigma) for Mouse 448 and human experiments or 1 mg/ml Hoechst (Sigma) for the mouse atlas experiments, followed by 5 washes with 2X SSC. The sample was covered with 24 × 32 mm coverslip spaced by a parafilm gasket and an overview image of the sample was taken using a 10X magnification objective. The area of the sample was traced on the overview image and used to generate fields of view (FOV) positions for the 40X and 60X objectives to cover the entire sample. Then the nuclei and Europium beads were imaged at 40X magnification and the FOV positions for the 60X objectives were saved together with the relative locations of the reference crosses for later alignment.

Afterwards, the tissue was permeabilized for 5 minutes with 0.1% (v/v) Triton X-100 (Sigma), 10 mM Dithiothreitol (DDT, Sigma) in 1X Tris-Borate-EDTA (TBE, Thermo), and washed 5 times with 1X TBE. An electrical wire was mounted on the ITO surface with a conductive copper foil tape (m.nu). The cathode was an uncoated ITO coverslip. The slide was mounted in the EEL holder (Extended Data Fig. 1c) and two 1.5-mm thick PDMS (Sylgard 184 Dow Corning) spacers were placed on either side of the sample and covered with the cathode slide. The electrophoresis buffer containing 10 mM DTT and 1 M urea (Sigma) in 1X TBE was injected with a gel-loading pipette tip, after which the wires were connected to a RND 320-KA3005D laboratory power supply or a Keithly 2450 sourcemeter, and a potential of 1.5V (10 V/cm) was applied for 20 minutes. However, a 1.5 V battery also works. To ensure hybridization, the sample was subsequently incubated for 5 minutes with a high salt concentration buffer (6X SSC, this step was not yet implemented for the mouse atlas experiments). The sample was then washed twice with 2X SSC and digested by 3 incubations (10 minutes for mouse and 5 min for human) with digestion buffer containing 1% (w/v) sodium-dodecyl-sulfate (SDS, Sigma), 20 mM Tris HCl (Thermo), 2,000 U/ml Superase (Thermo) and Proteinase K (1 U/ml for mouse and 0.5 U/ml for human) of at 30°C. Followed by 3 washes of 5 minutes with warm 5% (w/v) SDS in 2X SSC at 30 °C and 5 washes with 2X SSC at RT.

For non-barcoded experiments, the captured RNA was detected with directly labelled smFISH probes (Biosearch tech) diluted to 250 nM per probe in hybridization mix containing 10% (v/v) deionized formamide (Ambion), 0.1 g/ml dextran sulphate (Mw > 500,000, Sigma), 1 mg/ml *E. coli* tRNA (Roche), 2 mM Ribonucleoside Vanadyl Complexes (RVC, Sigma) and 200 μg/ml Bovine Serum Albumin (BSA, Sigma) in 2X SSC for at least 30 minutes at 38.5°C. Unbound probes were washed away with 3 washes of 10 minutes with 20% formamide in 2X SSC at 38.5 °C and 5 washes with 2X SSC. The sample was subsequently mounted on a microscope slide with Prolong Glass Antifade mounting media (Thermo).

For barcoded experiments, the RNA was fixed on the surface for 10 minutes with 4% PFA (Sigma) in 1X PBS (Thermo), followed by 5 washes with 2X SSC. The appropriate amount of encoding probes was dried using a SpeedVac depending on the probe production’s final concentration as measured by Qubit (Thermo) and resuspended in 20 μl of hybridization mix with 30% formamide. The mix was then carefully pipetted on the sample, covered with a plastic Hybrislip, placed in a petri dish humidified with SSC 2X and incubated for at least 24 hours at 38.5°C. Afterwards, the Hybrislip was carefully removed by first adding some 2X SSC to the edge until it was absorbed under the Hybrislip creating space between slide and Hybrislip.

The slide was then mounted in a flow cell (Rebus Biosystems) and placed in the ROBOFISH fluidic system, which automatically performed all subsequent steps. The sample was flushed with 2X SSC and washed 4 times 15 minutes with 30% formamide in 2X SSC at 47 °C, followed by 4 washes with 2X SSC. The encoding probes were then fixed for 10 minutes with 10% PFA (Sigma) in 1X Phosphate buffered saline (PBS) and washed with 2X SSC. Fluorescent detection probes were dispensed to the sample at a concentration of 50 nM in 10% formamide hybridization mix and hybridised for 10 minutes at 37°C, followed by 3 washes of 3 minutes with 20% formamide in 2X SSC and 4 washes with 2X SSC. Imaging buffer containing 2 mM 6-hydroxy-2,5,7,8-tetramethylchroman-2-carboxylic acid (Trolox, Sigma), 5mM 3,4-Dihydroxybenzoic acid (DBA, Sigma) and 20 nM Protocatechuate 3,4-Dioxygenase from *Pseudomonas sp*. (PCD, Sigma) was then dispensed to the flow cell. Imaging was triggered by the ROBOFISH system and performed on a Nikon Ti2 microscope at 20 °C^40^. After imaging, the sample was washed 4 times with 2X SSC and fluorophores were cleaved off by reducing the thiol bond with 50 mM tris(2-carboxyethyl)phosphine (TCEP, Sigma) in 2X SSC during two washes of 10 minutes at 22 °C. This was followed by 10 washes with 2X SSC. Then 15 cycles of hybridization, washing, imaging and stripping were performed to image all 16 bits of the barcode.

Usually, two staggered experiments were run in parallel. While the first experiment was being imaged (2-3 days), the next experiment was started, hence reducing the down-time of the microscope and doubling the speed by which datasets were generated.

### ROBOFISH automated fluidics

The ROBOFISH system is an open-source fully automated fluidics and temperature control platform integrated with imaging. It is designed to dispense arbitrarily small volumes to a flow cell by bridging the dead volume so that costly solutions like probe mixes are not wasted. It is designed to be flexible, both in terms of components and running protocols. A syringe pump (Tecan Cavro XE 1000 or Cavro XCalibur) with a Y-valve is connected to the running buffer (2X SSC) and a reservoir tube. The reservoir tube is connected to two 10-port actuated valves (MX-II IDEX) which are in turn linked to buffer tubes, up to two flow cells and a waste container.

This setup enables the aspiration of the target buffer into the reservoir, after which extra running buffer is aspirated into the syringe pump so that, when dispensed, the extra volume bridges the dead volume between valve and flow cell (Extended Data Fig. 1e). Between the valve and flow cell, a bubble-trap (Elveflow) and liquid degasser (Degasi Biotech) ensure that no air enters the flow cell. The flow cell itself can either be a flow cell designed by Rebus Biosystems, which is temperature controlled by a TC-720 controller (TE Technology Inc.), or the FCS2 flow cell from Bioptechs, that can either be temperature controlled by the Solid State Oasis or the Solid State ThermoCube recirculating chillers. The FCS2 flow cell temperature monitoring is implemented with the Yoctopuce Thermistor. Open-source Python drivers are available (TC-720: https://github.com/linnarsson-lab/Py_TC-720, ThermoCube: https://github.com/linnarsson-lab/ThermoCube, Oasis: https://github.com/linnarsson-lab/Oasis_chiller, MXII: https://github.com/linnarsson-lab/MXII-valve).

The ROBOFISH system monitors buffers volumes and notifies the user via text messages (Pushbullet, Pushbullet python API) when buffers need to be replaced or in the unlikely case of system errors or abnormal temperatures in the room. The full protocol is written to log- and metadata-files to save all information with the image datasets. Full building instructions, code and operating instructions are available online at https://www.protocols.io/view/robofish-construction-bcrciv2w and https://github.com/linnarsson-lab/ROBOFISH.

### Imaging

Imaging was performed on a Nikon Ti2 epifluorescence microscope equipped with a Nikon CFI Plan Apo Lambda 60X oil immersion objective with an numerical aperture (NA) of 1.4, CFI Plan Apo Lambda 40X objective with NA 0.95, CFI Plan Apo Lambda with NA 0.45, Nikon motorised stage, Nikon Perfect Focus system, Sona 4.2B-11 back-illuminated sCMOS camera with 11um pixels (Andor), Lumencor Spectra light engine (for configuration see Table 1), matching filter sets (see Table 2).

**Table 1.** Light source specifications.

| Spectral name | Bandpass (nm) | Power (mW) |
| --- | --- | --- |
| violet | 360/28 | 400 |
| cyan | 494/20 | 400 |
| green | 534/20 | 400 |
| yellow | 586/20 | 400 |
| red | 631/28 | 500 |
| red 2 | 690/10 | 500 |
| NIR | 747/11 | 500 |
| NIR2 | 780/10 | 500 |

**Table 2.** Filter cube specifications.

| Name | Emission | Dichroic | Excitation |
| --- | --- | --- | --- |
| multiband-89403m-dapi-A590-A750 | 89403m<br>https://www.chroma.com/products/parts/89403m | 89403bs<br>https://www.chroma.com/products/parts/89403bs | 89403X chroma<br>https://www.chroma.com/products/parts/89403x |
| ET525/30-A488 | ET525/30m<br>https://www.chroma.com/products/parts/et525-30m | T505lpxr<br>https://www.chroma.com/products/parts/t505lpxr | None |
| ET575/40m-A532 | ET575/40m<br>https://www.chroma.com/products/parts/et575-40m | T550lpxr<br>https://www.chroma.com/products/parts/t550lpxr | None |
| ET667/30-A647 | ET667/30<br>https://www.chroma.com/products/parts/et667-30m | T647lpxr<br>https://www.chroma.com/products/parts/t647lpxr | None |
| ET740/40X-IR700 | ET740/40X<br>https://www.chroma.com/products/parts/et740-40x | ZT670rdc-xxrxt<br>https://www.chroma.com/products/parts/zt670rdc-xxrxt | None |
| RET792lp-IR800 | FF01-832/37-32<br>https://www.semrock.com/filterdetails.aspx?id=ff01-832/37-25 | RT785rdc<br>https://www.chroma.com/products/parts/rt785rdc | None |

The automated image acquisition protocol was made in Nikon NIS Elements as a custom Job (available at https://github.com/linnarsson-lab/ROBOFISH). Data acquisition is triggered by the ROBOFISH system and images of each FOV with a Z-stack of 17 slices with a 0.3 μm step. Acquisition of a Z-stack was necessary to compensate for the curvature of the coverslip, caused by the assembly procedure in the flow cell. A typical sagittal mouse brain section was covered by 500-700 FOV positioned with an 8% overlap. Alexa-647 and Europium were imaged using the ET667/30-A647 filter cube by switching between 631 nm and 360 nm excitation at 100% and 30% power respectively, and an exposure of 80 ms for both. After imaging completion, the imaging Job notified the ROBOFISH system to continue with the fluidics of the next cycle.

### Surface pH measurement

The surface pH during electrophoresis was measured by fluorescent readout of *E. coli* particles doped with pH sensitive pHrodo Green fluorophore (Thermo). 200 μl of 1 mg/ml sonicated particles in 1X PBS were deposited on the capture slide after Europium bead deposition (Extended data Fig. 2a,b). Afterwards, a 10 μm mouse brain tissue section was placed on top and permeabilized as described above. A series of buffers with varying pH (MES buffer: 4.7, 5.0, 5.5, 6.0, 6.5, SSC: 7.0 and PBS: 7.4) was used to make a fluorescence calibration curve by measuring the mean pixel intensities of the particles at different pH levels (494 nm excitation, 200 ms exposure time using the 60X objective) (Extended data Fig 3c). Large aggregates of particles were excluded from the analysis.

An additional sample was placed in the electrophoresis holder, electrophoresis buffer was injected (TBE pH 8.3, 1M Urea, without DTT) and mounted on the microscope. Images of 2 FOV were taken every 45 seconds and after 2.5 minutes of baseline acquisition, electrophoresis at 1.5V was performed for 20 minutes. During this time the mean measured fluorescence did slightly increase but never in the range of the calibration curve, indicating that the buffer maintains the pH (Extended data Fig. 2d). Contrary, when higher voltages (4V and above) were applied, we did observe a drop in pH at the surface as interpolated with the calibration curve (6V example in Extended data fig. 2e,f). The higher the voltage, the faster the pH drop and the effect was more pronounced at surfaces covered by the tissue, thus indicating that restricted diffusion augments the surface pH change (data not shown).

### Image analysis pipeline

We developed a processing pipeline to automatically detect and decode the EEL signal (https://github.com/linnarsson-lab/pysmFISH_auto, Extended data Fig. 7). Briefly, after parsing, filtering and counting, the detected spots are registered between hybridizations using the Europium fiducial beads. The barcodes are then identified using a nearest neighbour approach. In order to map the EEL signal (60X objective) to the nuclei images (40X objective), we first registered the Europium beads imaged with both objectives by using point cloud registration^41^, and the resulting transformation was applied to the EEL signal. Nuclei were segmented using Cellpose^29^ and the segmentation masks were expanded by 10 μm without overlapping. Detected signal dots were then assigned to cell labels if they fell inside a segmentation mask making use of a k-d tree algorithm. In order to process the large amount of data generated by each EEL experiment, the analysis is parallelized using Dask^42^.

### Optical density

Optical density simulations were performed by filling a 10 × 10 um area with an increasing number of randomly spaced points and determining how many could be resolved using Abbe’s diffraction limit (λ / 2NA) as the minimal distance. The simulation was repeated for various wavelengths of light and for objectives with different numerical apertures.

### Spatial analysis mouse samples

Lateral diffusion was investigated by identifying *Vip+* cells in the osmFISH and the 7 mouse atlas experiments through DBSCAN. Their centroid was calculated based on the location of the molecules, after which the probability of finding a molecule in concentric circles around the centroid was calculated and compared.

To facilitate the exploration and analysis of point-based spatial datasets that contain millions of molecules, we developed a Python package called FISHscale (https://github.com/linnarsson-lab/FISHscale), which efficiently handles large datasets by storing them on disk and relying on parallelized processing with Dask^42^. FISHscale can handle multiple datasets for analysis and 3D visualisation. The visualisation tool is based on Open3D^43^, which enables rapid visualisation of millions of molecules.

FISHscale implements a method to regionalize the tissue sample by binning the data in hexagonal bins, performing Principal Component Analysis (PCA) or Latent Dirichlet Allocation (LDA), and clustering the hexagons using Ward hierarchical clustering making use of the spatial information by using a connectivity matrix for the neighbouring hexagonal tiles. The resulting region labels can be ordered by Spectral Embedding to give transcriptionally similar regions a similar color. If multiple regionalized datasets share anatomical structures between them, these regions can be linked by correlating their mean expression.

To visualise the mixture of region identities, a Random Forest Classifier was trained on the hexagonally binned data with the region labels, and the probability of each region label was obtained by querying the classifier. These probabilities were then multiplied by the RGB values of original region colors, and summed to display gradients in the form of mixed colors.

Border strength was calculated by placing a grid of points over the sample and selecting all molecules within a 200 μm radius from each point. Each group of selected molecules was then split in half 12 times at different angles. For each angle, we calculated the total number of molecules in each half and measured the Euclidean distance between the counts. A large distance corresponds to a bigger difference in gene expression between the regions separated along a specific angle. This allows us to measure both the strength and the angle of the potential border (Fig. 4b). In the 3D rendered image (Fig. 4f, Blender https://www.blender.org/), the border strength is visualised as the height of the mixed region colors.

To align the hexagonally binned spatial datasets with a previously published single-cell RNAseq study of the mouse nervous system^1^, we used a generalised version of Tangram^30^ called Bonefight^31^ to calculate the probability distributions of the location of each of the 199 cell types that were identified in the mouse brain. The neurotransmitter identity of the various cell types was summed to give the probability of the spatial location of each neurotransmitter.

### Spatial analysis of human samples

The human data was segmented using an unsupervised graph-based approach^44^ in which the algorithm enforces that connected nodes should have similar embeddings, whereas randomly sampled pairs of nodes should have dissimilar embedding. We built a graph neural network of two layers with 24 hidden units, a rectified linear unit activation function between the layers, a pooling aggregation, and a differentiable group normalisation. The graph was built using the Deep Graph Library’s SAGEconv module (https://www.dgl.ai/). Each RNA molecule formed a node in the graph and any two molecules were connected by an edge if their distance was below 15 μm. During training, mini-batches of 512 nodes were generated, for each centre node a neighbourhood 1-hop and 2-hops away is subsampled (maximum 20 nodes for the first hop and 10 nodes second hop), each layer aggregates and updates the information of each node and its sampled neighbourhood. After training, an embedding was generated for every molecule, and k-means clustering was used to cluster molecules into distinct spatial domains. The genes enriched in each spatial domain were used for annotation.

---

**Supplementary Fig. 1.**
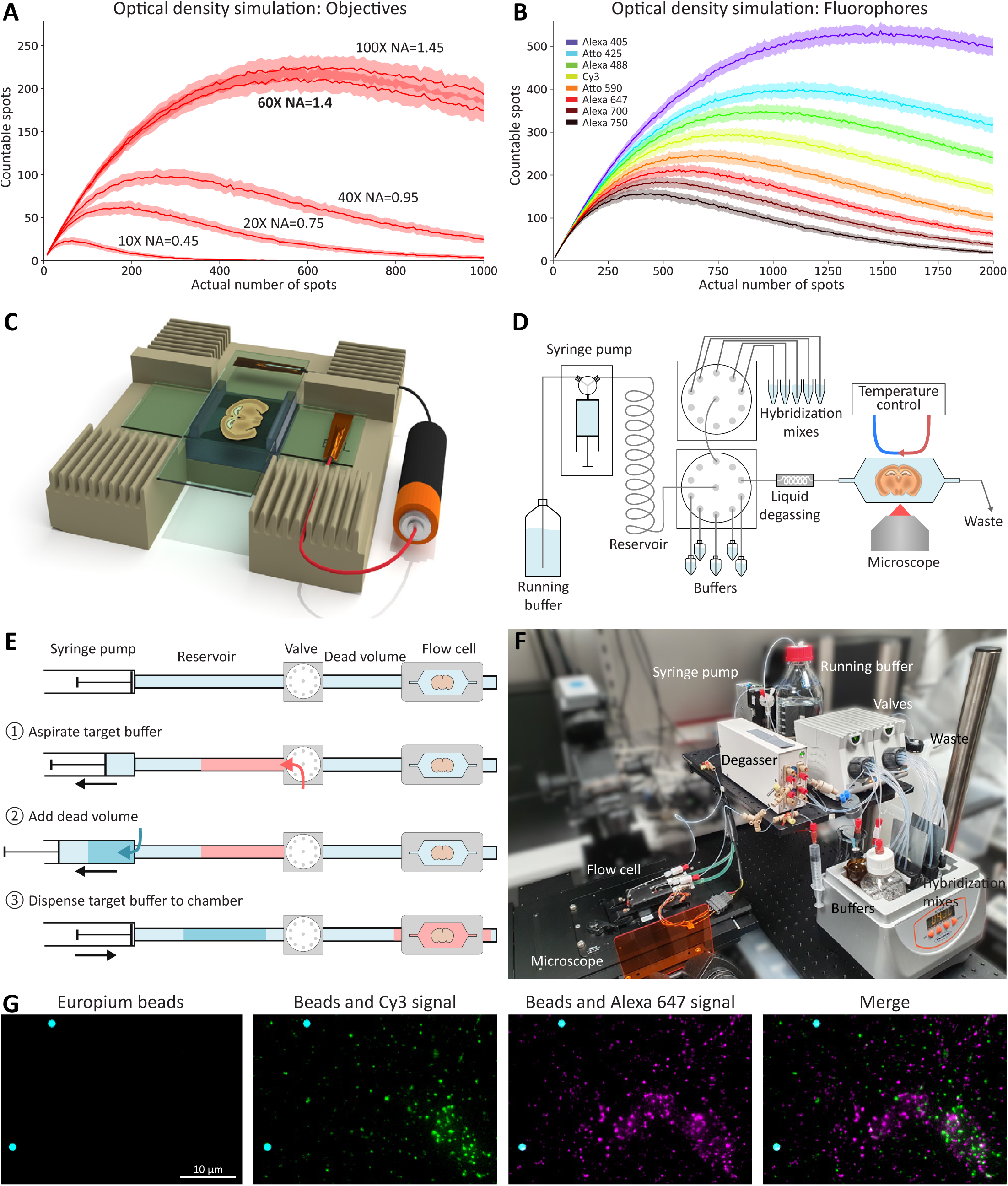
EEL setup. **a**, Computer simulation of countable dots, defined as not overlapping with any other dot within the Abbe diffraction limit, for increasing numbers of dots in a virtual cell for various objective lenses. **b**, Same simulation as in **a** but with the 60X NA=1.4 objective for various fluorophores. **c**, EEL electrophoresis setup. The tissue section on the capture slide is placed on the bottom of the 3D printed holder and connected to the positive pole of a power source. The top electrode is spaced with a silicone strip and connected to the negative pole of the power source. **d**, Simplified schematic of the ROBOFISH fluidic system. **e**, Working principle of the ROBOFISH fluidic system. The target liquid is aspirated into the reservoir. Then the dead volume is aspirated into the syringe. When the full volume is dispensed to the flow cell the target liquid will reach the flow cell, without having to fill the dead volume. **f**, Image of a ROBOFISH system. **g**, Example of two-color EEL experiment encoding up to 2 × 448 genes.

**Supplementary Fig. 2.**
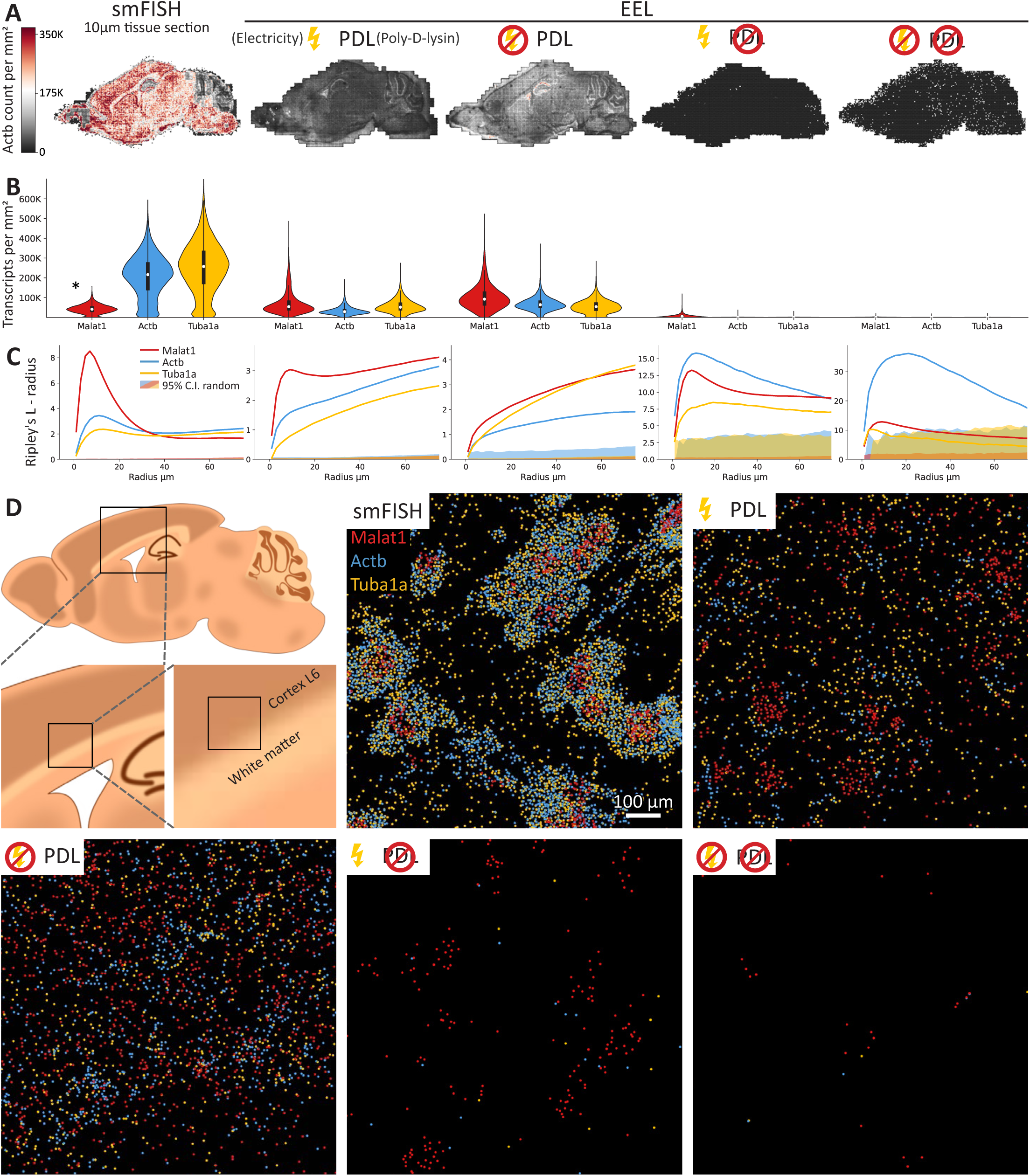
Effect of electricity and poly-D-lysine (PDL) on RNA transfer. **a**, Actb molecule counts in adjacent sections for: smFISH in a 10 μm tissue section, EEL with electrophoresis and PDL, EEL without electrophoresis but with PDL, EEL with electrophoresis but without PDL and EEL without electrophoresis and without PDL. Quantified in binned data and normalized to mm^2^. **b**, Quantification of 3 genes in the same experiments as in **a**, showing increased RNA capture in EEL conditions with PDL. * Malat1 is highly expressed, therefore the dots start to overlap and quantification is likely an underestimate of the real number of molecules. **c**, Ripley’s L score minus the radius for the same experiments as in **a**. Showing the 95% confidence interval (C.I., solid fill) of randomized data for the same amount of molecules, which center around 0, meaning no deviation from random. In the original situation (smFISH in tissue) the RNA locations of all genes show various degrees of clusterdness with a peak around the average cell size. The addition of electricity in the EEL protocol improves the clusterdness as compared to RNA transfer only by diffusion. **d**, Raw RNA dots for the same experiments and similar area, show a more cell-like RNA pattern for EEL conditions where electricity was used.

**Supplementary Fig. 3.**
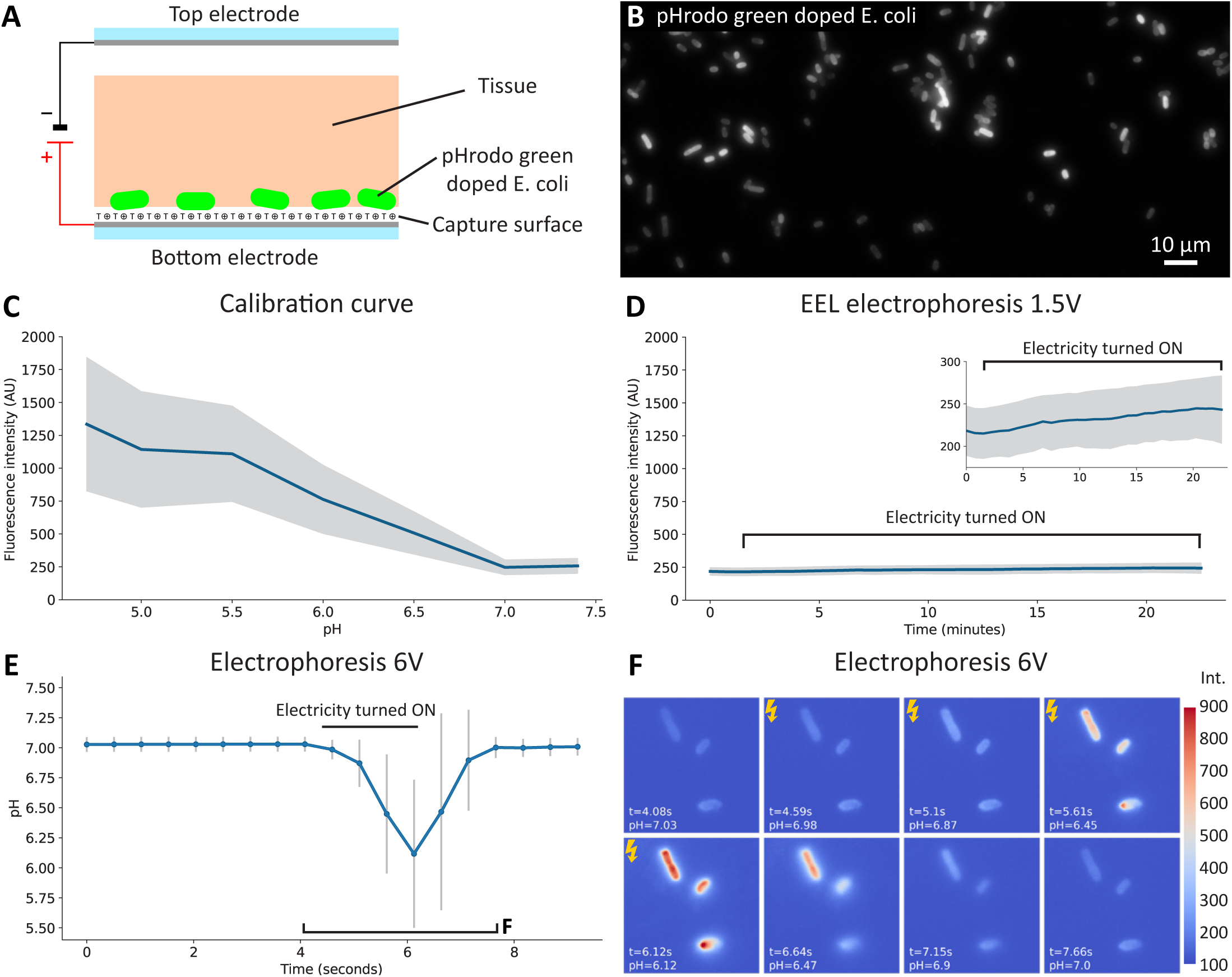
The EEL electrophoresis step does not lower surface pH. **a**, Experimental setup with pHrodo green doped E. coli sandwiched between the capture slide and the tissue, which is the place where the RNA will be captured and pH could have an effect. **b**, Image of particles at physiological pH. **c**, Calibration curve by measuring fluorescence intensity at various pH levels. **d**, Fluorescence intensity measured every 45 seconds during a regular 20 minute EEL electrophoresis step at 1.5 Volt. Data shown with the same y-scale as in **c**. A very slight increase was observed (see inset with rescaled y-axis). However, fluorescence intensity stayed below the calibration curve indicating no detrimental drop in pH. **e, f**, When performing electrophoresis at higher potentials, the surface pH rapidly drops.

**Supplementary Fig. 4.**
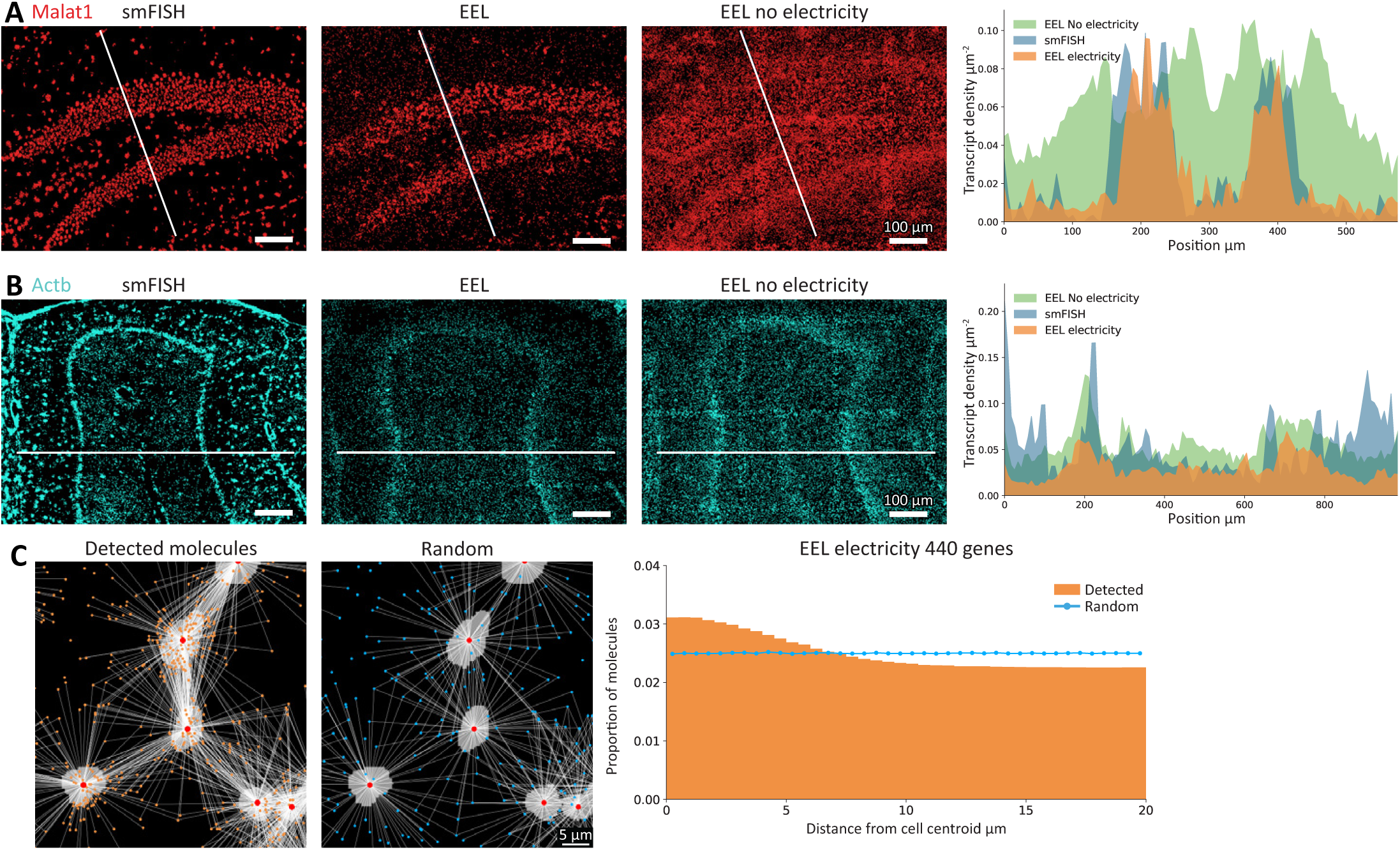
Transcript localization in EEL. **a, b**, Malat1 and Actb transcript localization in dentate gyrus and cerebellum respectively, for smFISH in a 10 μm section, EEL with electrophoresis and EEL without electrophoresis. Density profiles on the right show that EEL with electricity better matches the original tissue sturcture. In the case of Malat1 the no-electricity condition shows a very distorted blot. **c**, Measured distances between all detected molecules in the 440 genes sagittal mouse brain section and the nucleus centroid of the cells. Repeating this measurement with randomized data indicated that molecules are more likely to be found close to the centroids, supporting that the RNA transfer matches the sparse cellular architecture of the mouse brain.

**Supplementary Fig. 5.**
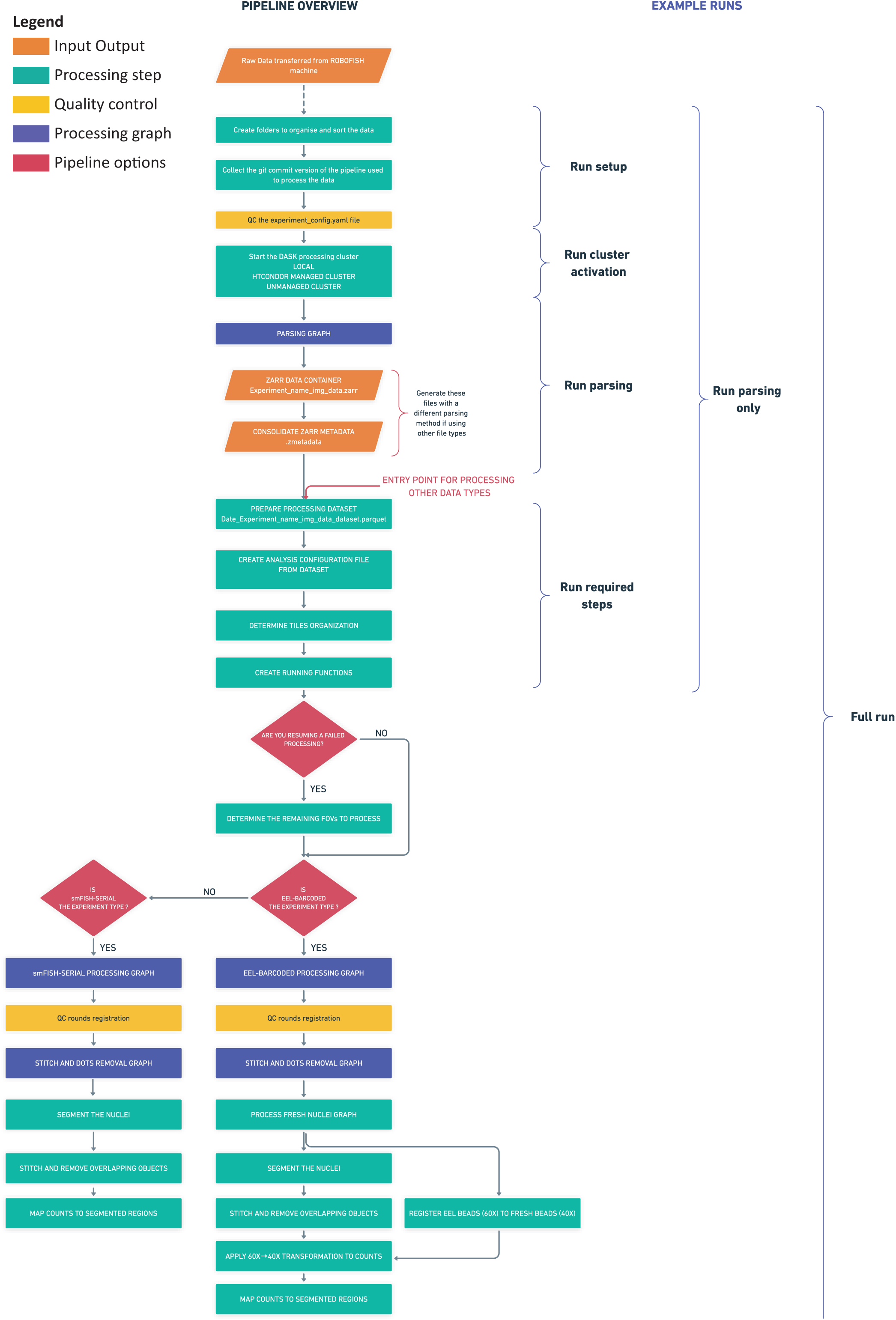
Image analysis pipeline of pysmFISH.

**Supplementary Fig. 6.**
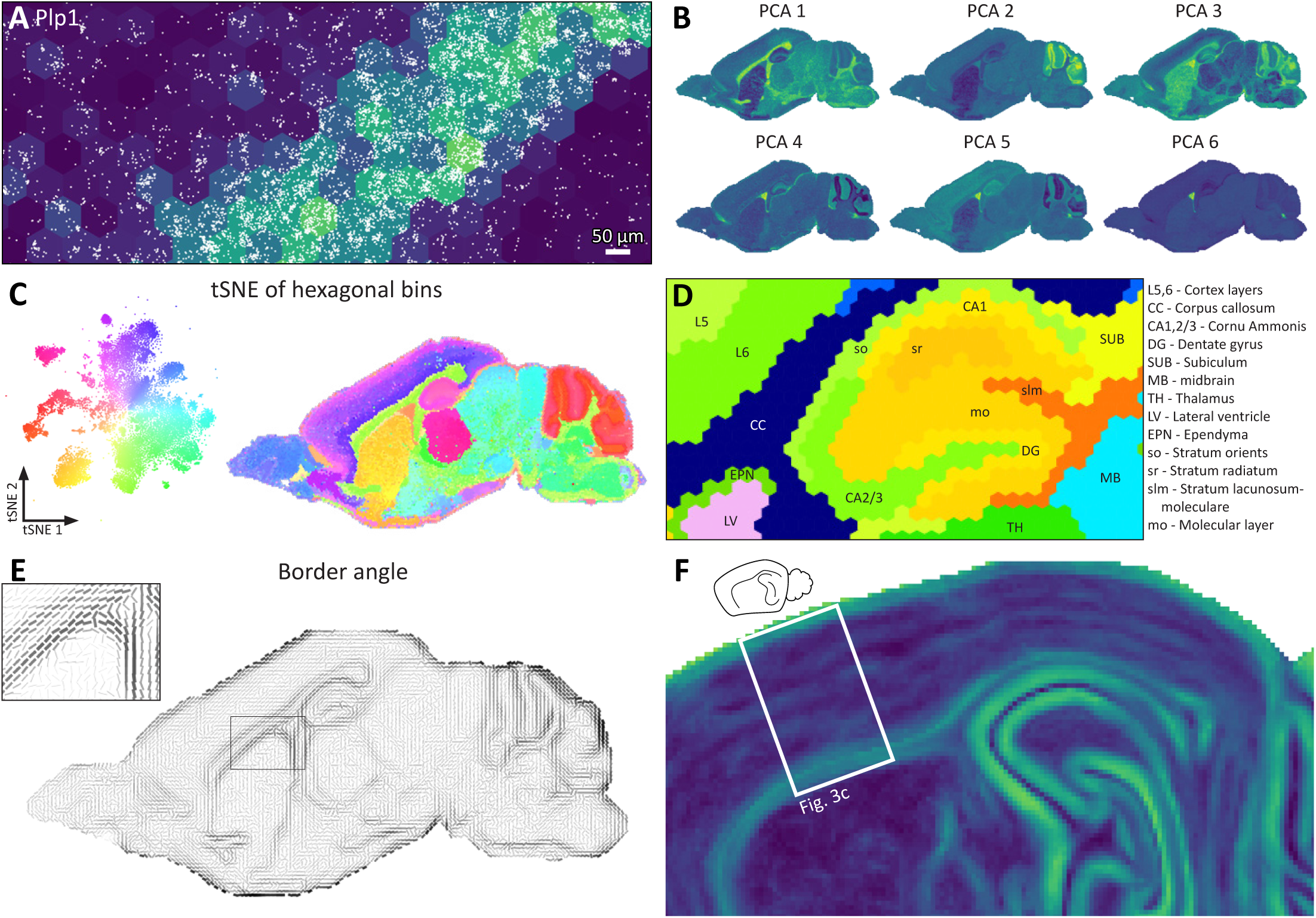
Data-driven regionalization of mouse brain. **a**, Hexagonal binning of detected signal, here showing Plp1 as an example. **b**, First 6 PCA components of hexagonally binned data shows that components capture different anatomical structures. **c**, t-SNE embedding of hexagonal bins also show that anatomical structures are captured by the components. **d**, Detailed view of the regionalization in the hippocampus with labelled anatomical regions. **e**, Angle of the largest transcriptional difference indicating border direction. Line width and darkness correspond to border strength. **f**, Border strength of section 6 of the mouse atlas that is used for Fig. 3c, showing borders in the cortex.

**Supplementary Fig. 7.**
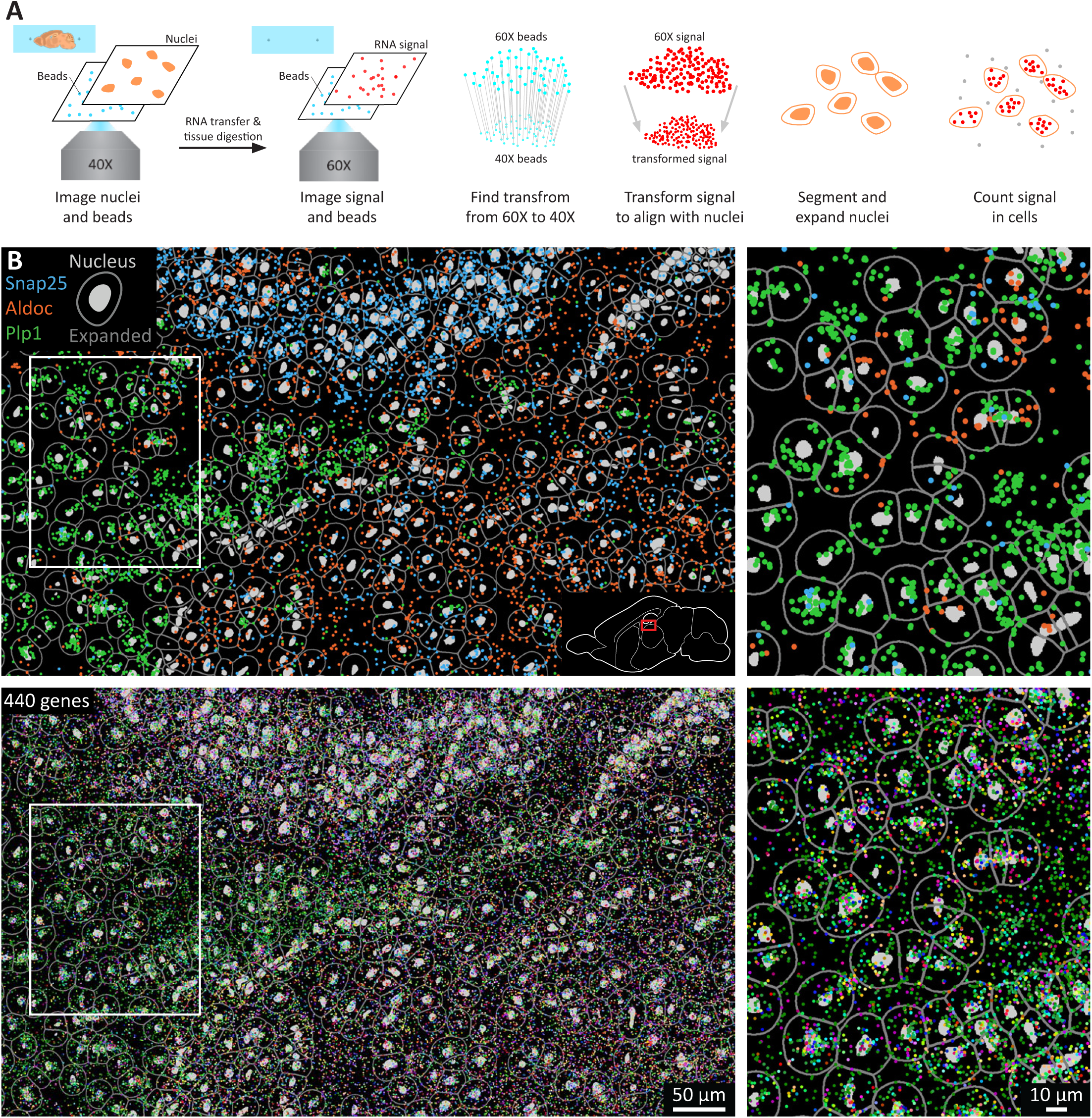
Single cell segmentation and counting. **a**, Pipeline to segment cells. Before the RNA is transferred and the tissue is digested, the nuclei are imaged at 40X magnification along with fiducial beads. These beads are also imaged when imaging the signal at 60X magnification so that they can be used to match the images before and after tissue removal. The signal is transformed to fit the space of the 40X images. Nuclei are segmented and expanded as a proxy for the cell boundary. Then, RNA is counted in the segmented and expanded masks to generate a gene-by-cell matrix. **b**, Example region showing segmentation in white matter, hippocampus and thalamus for 3 genes (top), or all 440 genes (bottom).

**Supplementary Fig. 8.**
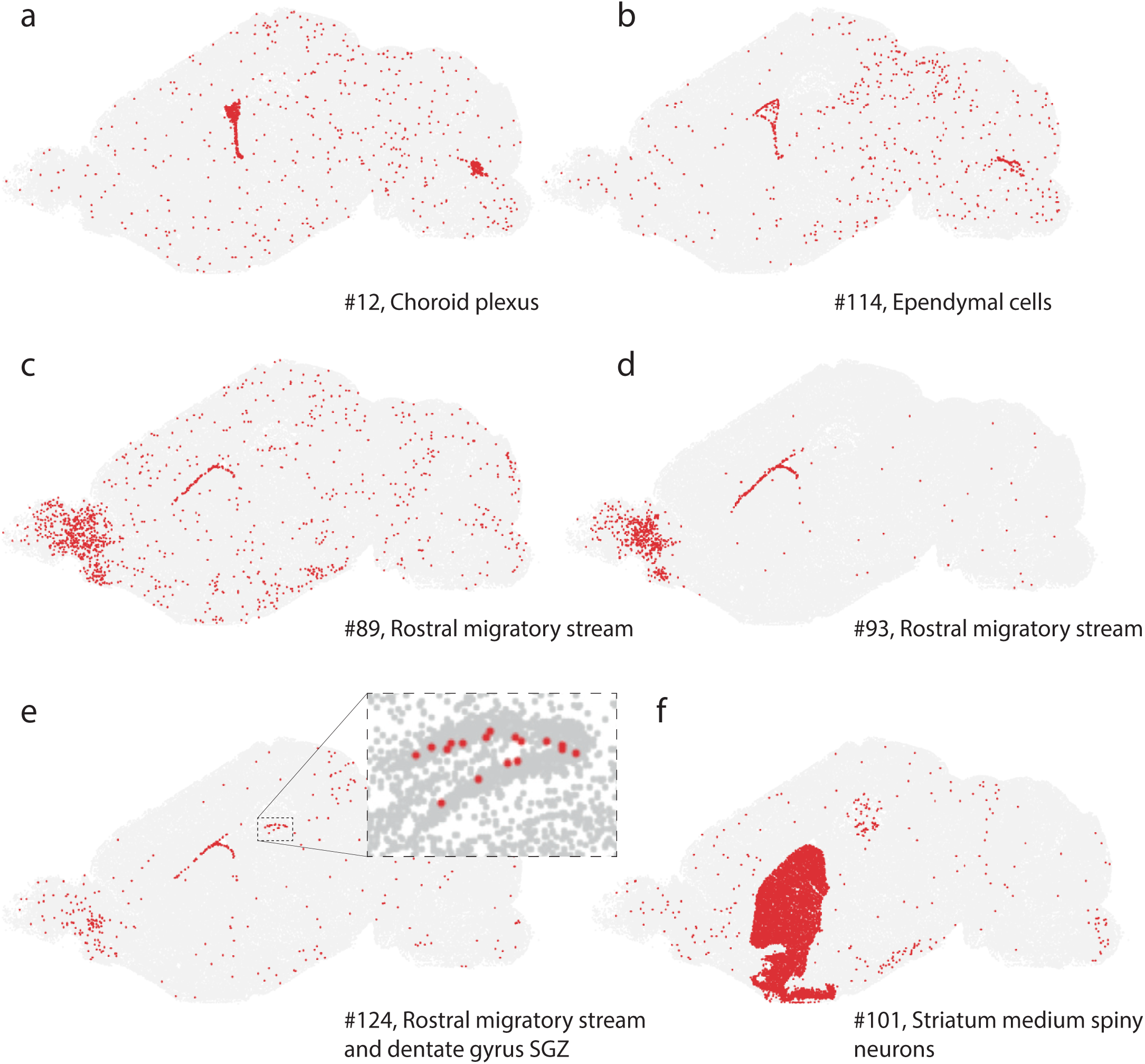
Clusters related to neurogenesis and the lateral ventricles. (**a**) Choroid plexus, expressing Kl, Foxj1, Aqp1. (**b**) Ependymal cells (Ccdc153, Foxj1, Tmem212). (c, d) Rostral migratory stream (Dlx1, Meis2, Sox11). (**e**) Subventricular zone of the lateral ventricle and subgranular zone of the dentate gyrus (Sox11, Igfbpl1, Hes5); inset confirms the location of dentate gyrus stem cells along the hilus border. (**f**) Striatum medium spiny neurons (Adora2a, Gpr88, Drd1, Drd2).

**Supplementary Fig. 9.**
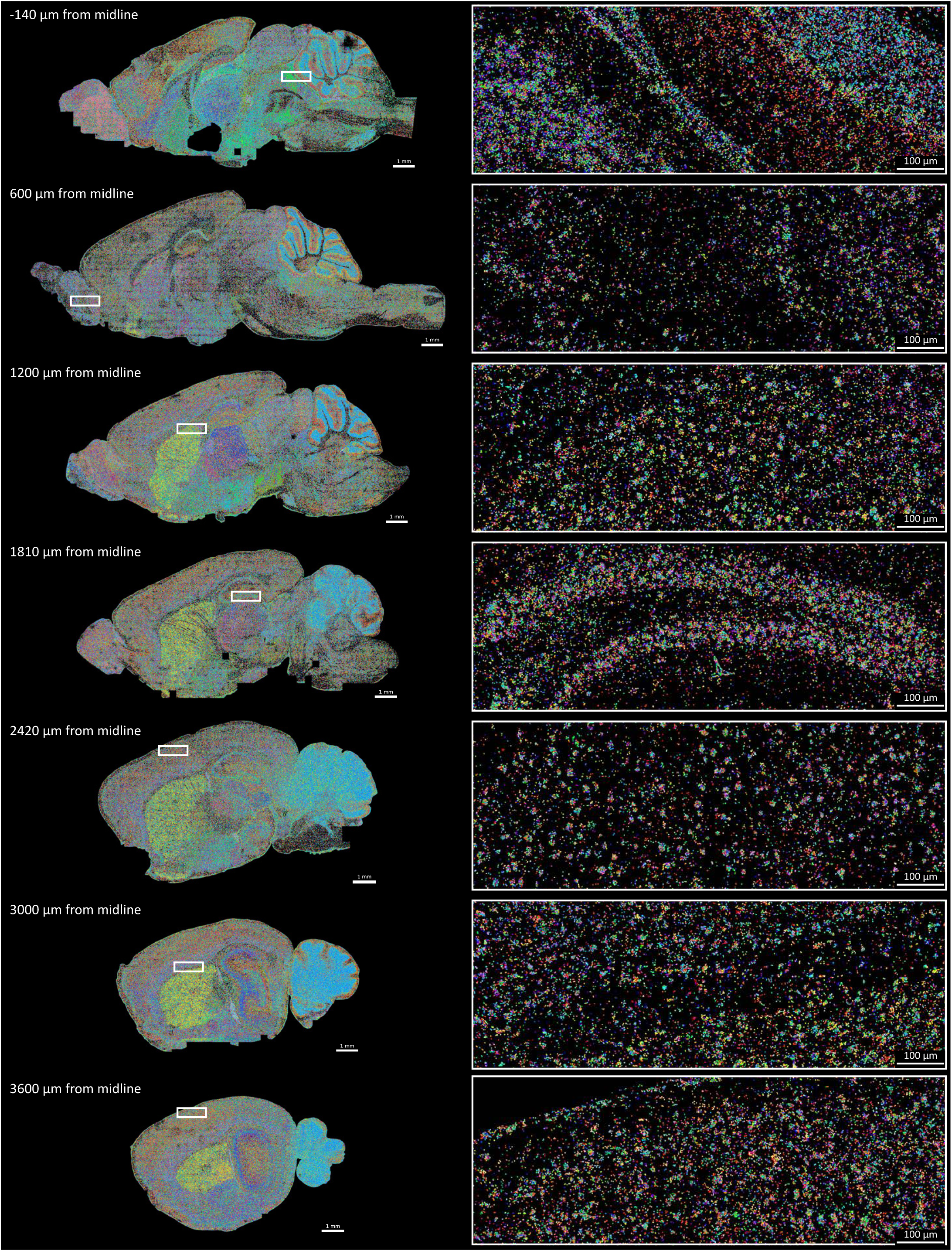
Raw data of the 7 sections of the sagittal mouse atlas. Colors correspond to one of 168 measured genes.

**Supplementary Fig. 10.**
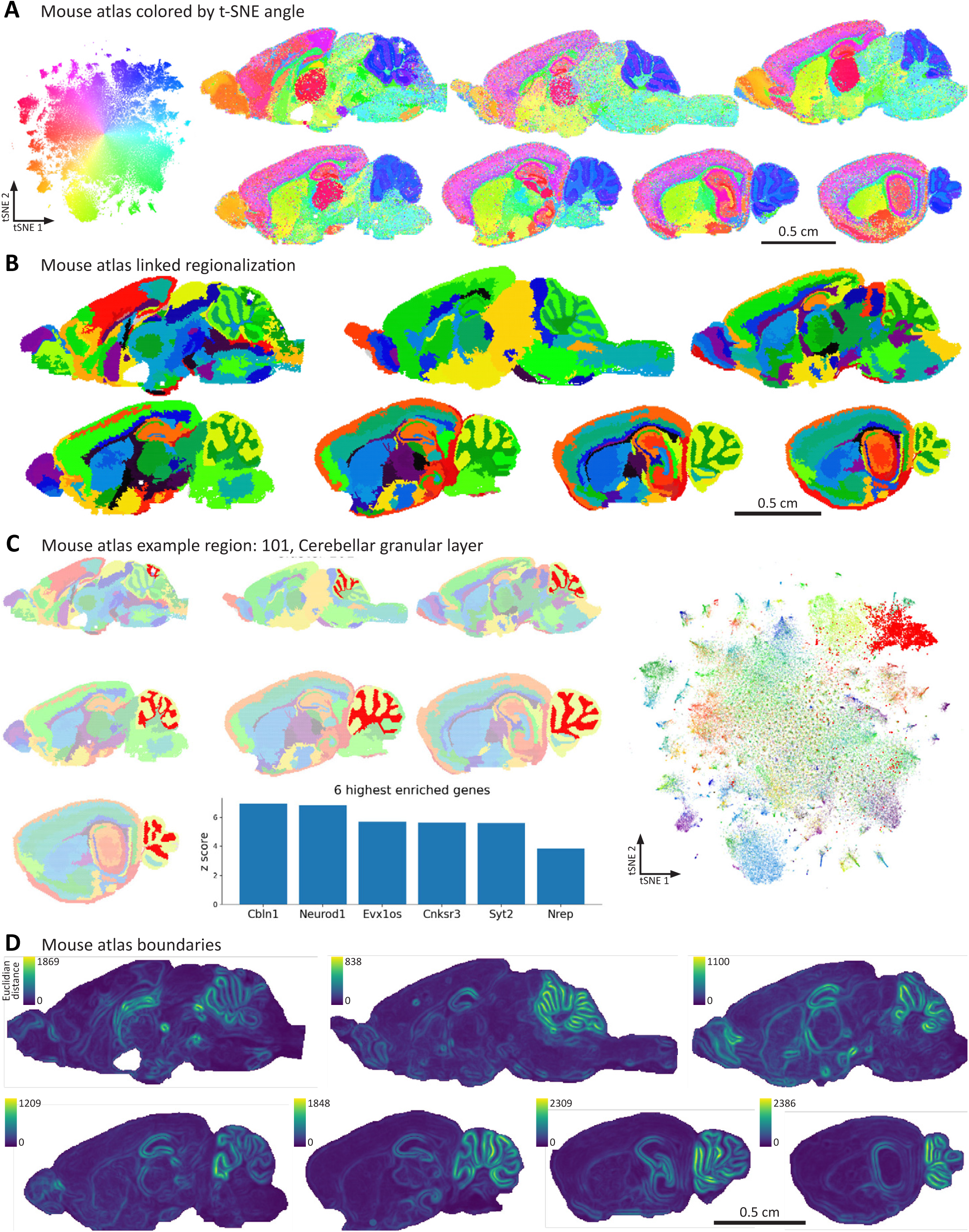
Mouse atlas regionalization and borders. **a**, t-SNE run on the combination of all hexagonal bins of the 7 sections, indicating that datasets integrate well as similar anatomical structures from adjacent sections co-localize on the t-SNE without showing obvious batch-to-batch variation. **b**, Regionalized mouse brain atlas where similar regions are linked between adjacent sections. **c**, Example of regions that link between all sections of the mouse atlas. Spatial location and location in the t-SNE is indicated in red. **d**, Boundary strength was measured on all sections and showed matching border locations between the same anatomical structures from adjacent sections.

**Supplementary Fig. 11.**
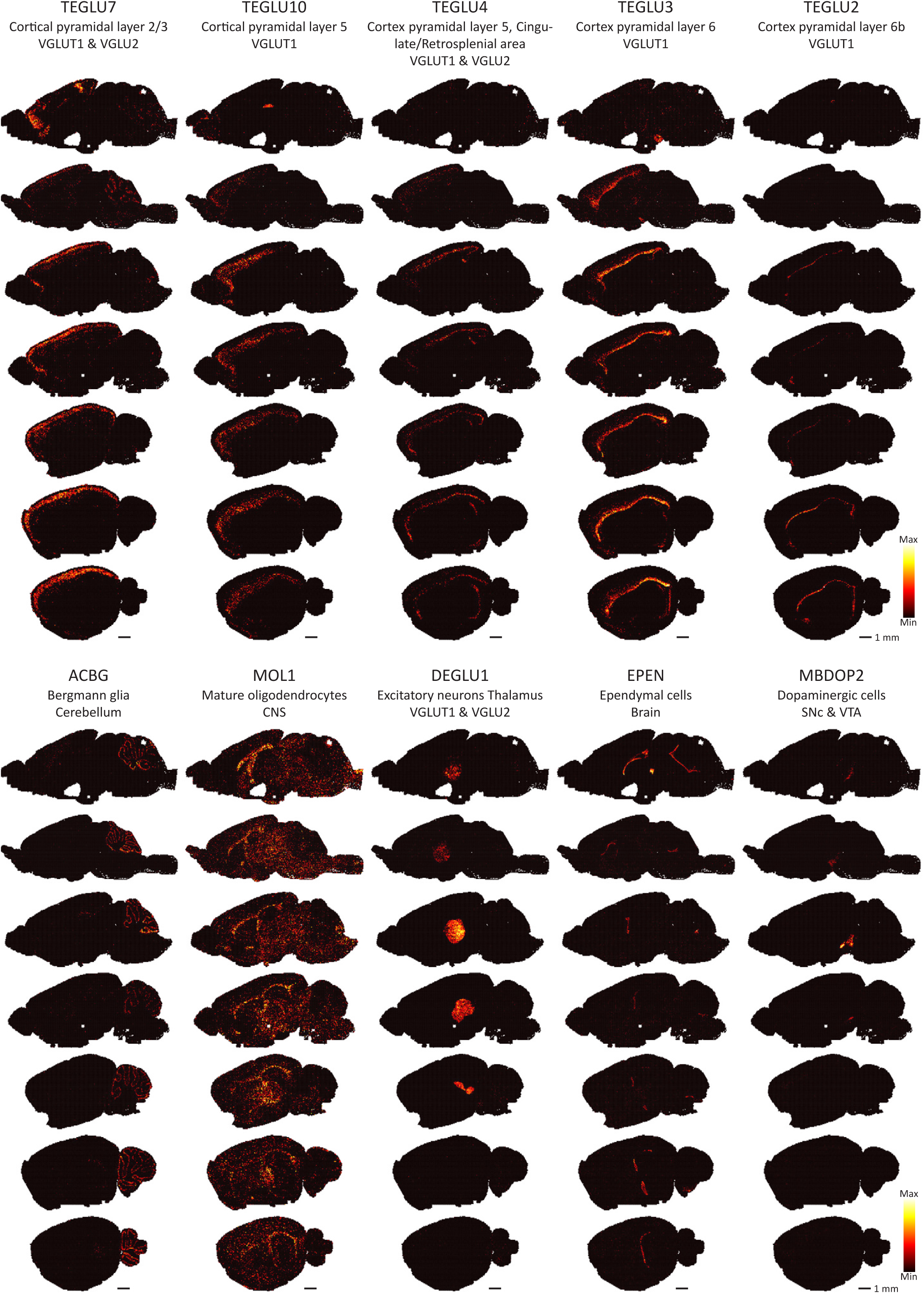
Integration of the spatial mouse atlas with single-cell RNAseq of the mouse brain. Likelihood of the spatial location of cell types as found by single-cell RNAseq.

**Supplementary Fig. 12.**
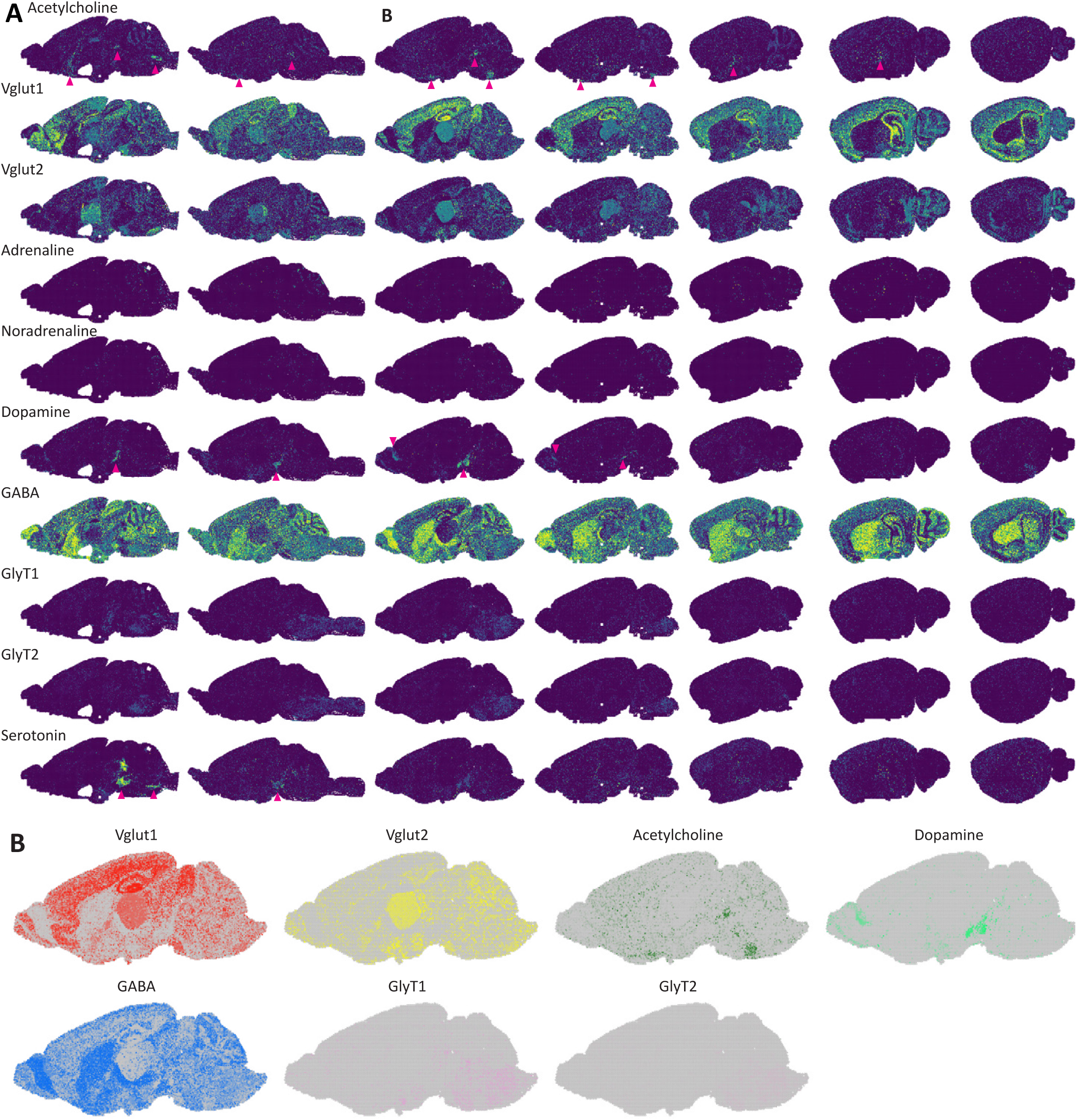
Integration of single-cell RNAseq data to find neurotransmitter domains. **a**, Likelihood of spatial location of neurotransmitters in the mouse brain atlas. Arrows indicate small nuclei of corresponding neurotransmitters. **b**, Integration of the location of 7 neurotransmitters in the 3rd section by mixing the colors of individual neurotransmitters shows both separated and shared neurotransmitter domains.

**Supplementary Fig. 13.**
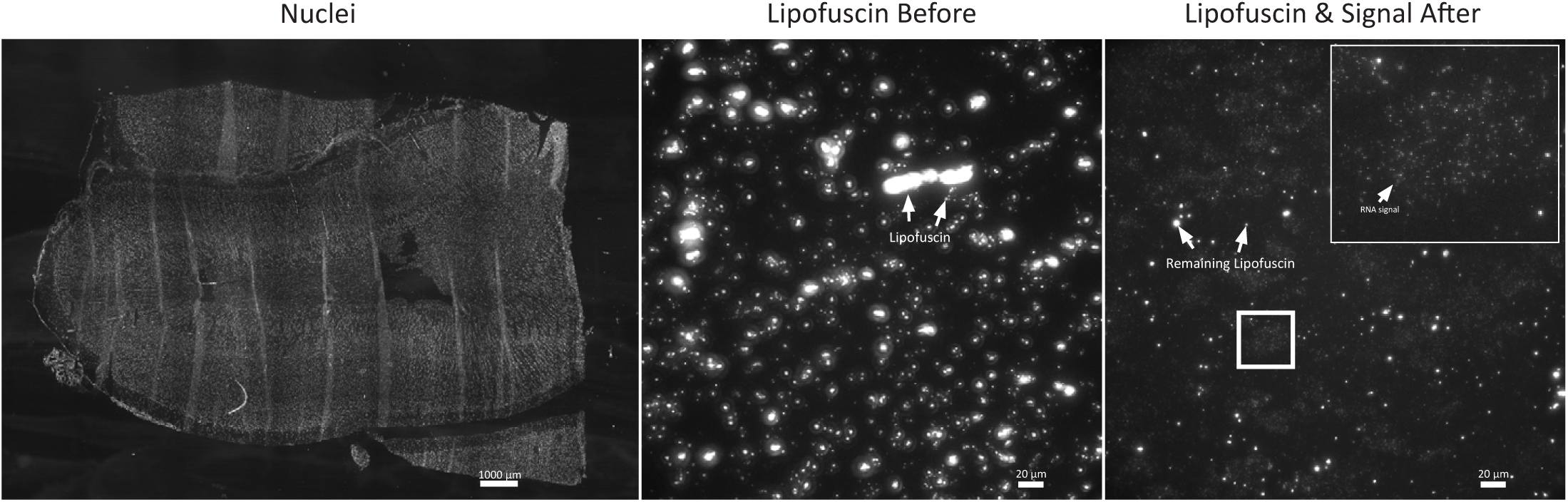
EEL significantly reduces lipofuscin. The left panel shows the nuclei image of the human adult primary visual cortex of Fig. 6 rotated 90 degrees to the left. Images before and after RNA transfer and tissue digestion show a stark reduction in lipofuscin content. Small dots on the right image correspond to the RNA signal spots, while brighter dots are remaining lipofuscin. Images were rescaled to the same minimum and maximum, and were taken with the same objective, illumination settings and exposure time.

**Supplementary Fig. 14.**
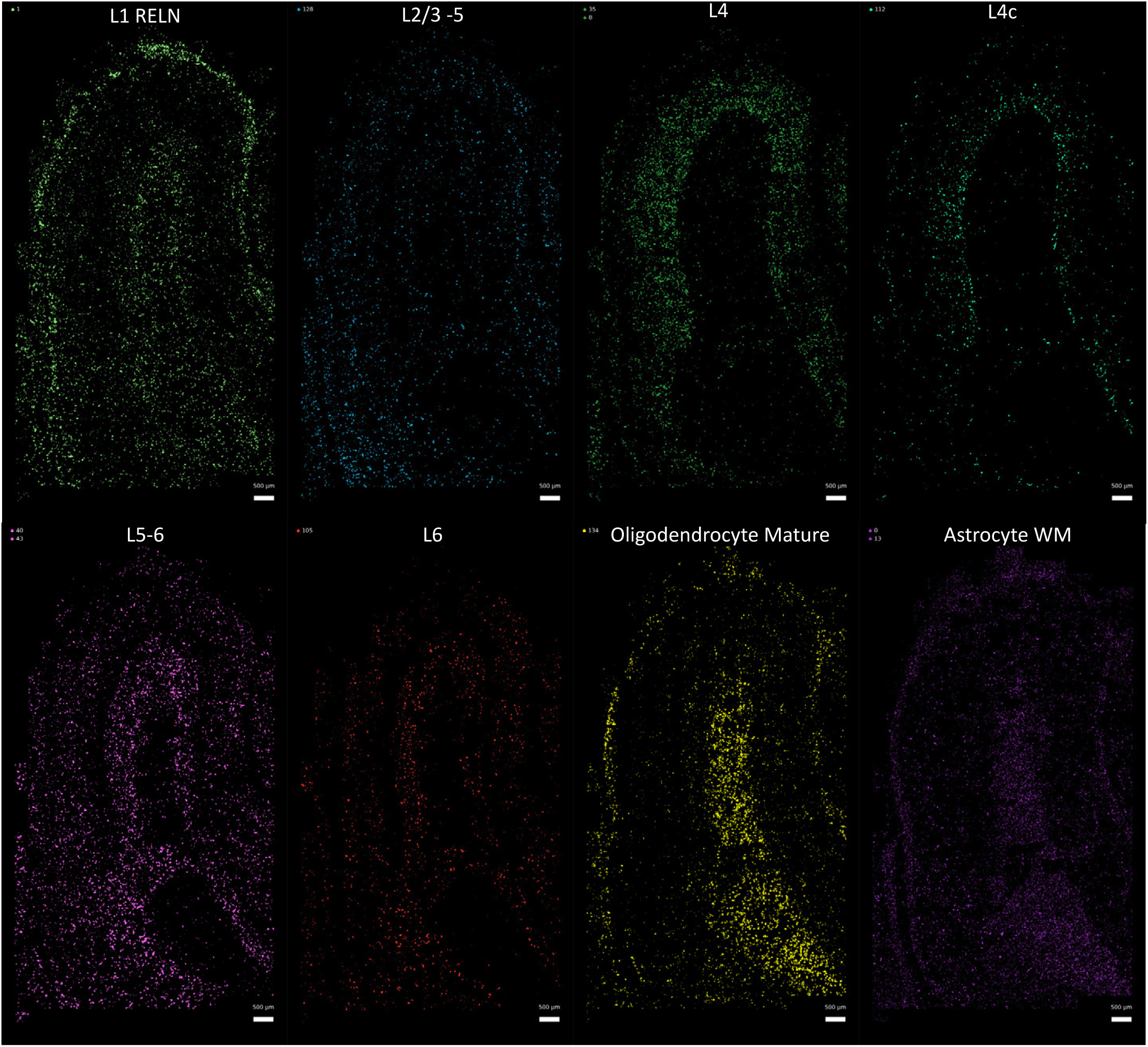
Application of GraphSAGE to the human visual cortex. Cell types linked to some of the GraphSAGE identified clusters after molecule embedding. Inh L1: Layer 1 inhibitory neurons, Exc L2/3-5: Layer 2/3 and 5 excitatory neurons, Exc L4: Layer 4 excitatory neurons, Exc L4c: Layer 4c excitatory neurons, Exc L5-6: Layer 5 and 6 excitatory neurons, Exc L6: Layer 6 excitatory neurons, Oligo: Mature Oligodendrocytes, Astro W.M: white matter Astrocytes and Astro L1-L6: astrocytes in layers 1 to 6.

## Notes

http://mousebrain.org

https://www.protocols.io/view/eel-fish-t92er8e

https://www.protocols.io/view/robofish-construction-bcrciv2w

https://github.com/linnarsson-lab/ROBOFISH

https://github.com/linnarsson-lab/FISHscale

https://github.com/linnarsson-lab/pysmFISH_auto

https://github.com/linnarsson-lab/oligopy

## References

1. Zeisel, A. et al. Molecular Architecture of the Mouse Nervous System. Cell 174, 999-1014.e22 (2018).

2. Codeluppi, S. et al. Spatial organization of the somatosensory cortex revealed by osmFISH. Nature Methods 15, 932–935 (2018).

3. Moffitt, J. R. et al. Molecular, spatial and functional single-cell profiling of the hypothalamic preoptic region. Science 5324, eaau5324 (2018).

4. Eng, C. H. L. et al. Transcriptome-scale super-resolved imaging in tissues by RNA seqFISH+. Nature (2019) doi:10.1038/s41586-019-1049-y.

5. Gyllborg, D. et al. Hybridization-based in situ sequencing (HybISS) for spatially resolved transcriptomics in human and mouse brain tissue. Nucleic Acids Research 48, E112 (2020).

6. Ke, R. et al. In situ sequencing for RNA analysis in preserved tissue and cells. Nature methods 10, 857–60 (2013).

7. Moses, L. & Pachter, L. Museum of Spatial Transcriptomics. bioRxiv (2021) doi:https://doi.org/10.1101/2021.05.11.443152.

8. Lein, E., Borm, L. E. & Linnarsson, S. The promise of spatial transcriptomics for neuroscience in the era of molecular cell typing. Science 358, 64–69 (2017).

9. Stahl, P. L. et al. Visualization and analysis of gene expression in tissue sections by spatial transcriptomics. Science 353, 78–82 (2016).

10. Vickovic, S. et al. High-definition spatial transcriptomics for in situ tissue profiling. Nature Methods 16, 987–990 (2019).

11. Rodriques, S. G. et al. Slide-seq: A scalable technology for measuring genome-wide expression at high spatial resolution. Science 363, 1463–1467 (2019).

12. Stickels, R. R. et al. Highly sensitive spatial transcriptomics at near-cellular resolution with Slide-seqV2. Nature Biotechnology 39, 313–319 (2021).

13. Fu, X. et al. Continuous Polony Gels for Tissue Mapping with High Resolution and RNA Capture Efficiency. bioRxiv 2021.03.17.435795 (2021).

14. Chen, A. et al. Title: Large field of view-spatially resolved transcriptomics at nanoscale resolution Short title: DNA nanoball stereo-sequencing. bioRxiv 2021.01.17.427004 (2021).

15. Cho, C.-S. et al. Seq-Scope: Submicrometer-resolution spatial transcriptomics for single cell and subcellular studies. bioRxiv 2021.01.25.427807 (2021).

16. Raj, A., van den Bogaard, P., Rifkin, S. a, van Oudenaarden, A. & Tyagi, S. Imaging individual mRNA molecules using multiple singly labeled probes. Nature methods 5, 877–879 (2008).

17. Femino, A. M., Fay, F. S., Fogarty, K. & Singer, R. H. Visualization of single RNA transcripts in situ. Science (New York, N.Y.) 280, 585–590 (1998).

18. Chen, K. H., Boettiger, a. N., Moffitt, J. R., Wang, S. & Zhuang, X. Spatially resolved, highly multiplexed RNA profiling in single cells. Science 348, aaa6090–aaa6090 (2015).

19. Lee, J. H. et al. Highly multiplexed subcellular RNA sequencing in situ. Science (New York, N.Y.) 343, 1360–3 (2014).

20. Shah, S., Lubeck, E., Zhou, W. & Cai, L. In Situ Transcription Profiling of Single Cells Reveals Spatial Organization of Cells in the Mouse Hippocampus. Neuron 92, 342–357 (2016).

21. Wang, X. et al. Three-dimensional intact-tissue sequencing of single-cell transcriptional states. Science 361, eaat5691 (2018).

22. Xia, C., Fan, J., Emanuel, G., Hao, J. & Zhuang, X. Spatial transcriptome profiling by MERFISH reveals subcellular RNA compartmentalization and cell cycle-dependent gene expression. Proceedings of the National Academy of Sciences 116, 201912459 (2019).

23. Ripley, B. D. The second-order analysis of stationary point processes. Journal of Applied Probability 13, 255–266 (1976).

24. Ortiz, C. et al. Molecular atlas of the adult mouse brain. Science Advances 6, 1–14 (2020).

25. Partel, G. et al. Identification of spatial compartments in tissue from in situ sequencing data. BMC Biology 18, 144 (2020).

26. Stanley, G., Gokce, O., Malenka, R. C., Südhof, T. C. & Quake, S. R. Continuous and Discrete Neuron Types of the Adult Murine Striatum. Neuron 105, 688-699.e8 (2020).

27. Petukhov, V. et al. Cell segmentation in imaging-based spatial transcriptomics. Nature Biotechnology (2021) doi:10.1038/s41587-021-01044-w.

28. Park, J. et al. Cell segmentation-free inference of cell types from in situ transcriptomics data. Nature Communications 12, 1–13 (2021).

29. Stringer, C., Wang, T., Michaelos, M. & Pachitariu, M. Cellpose: a generalist algorithm for cellular segmentation. Nature Methods 18, 100–106 (2021).

30. Biancalani, T. et al. Deep learning and alignment of spatially resolved single-cell transcriptomes with Tangram. Nature Methods 18, 1352–1362 (2021).

31. la Manno, G. et al. Molecular architecture of the developing mouse brain. Nature 596, 92–96 (2021).

32. Partel, G. & Wählby, C. Spage2vec: Unsupervised representation of localized spatial gene expression signatures. The FEBS Journal 288, 1859–1870 (2021).

33. Gennari, F. De Peculiari Structura Cerebri. (Ex Regio Typographeo, 1782).

34. Eng, C.-H. L., Shah, S., Thomassie, J. & Cai, L. Profiling the transcriptome with RNA SPOTs. Nature Methods 14, 1153–1155 (2017).

35. Lein, E. S. et al. Genome-wide atlas of gene expression in the adult mouse brain. Nature 445, 168–176 (2007).

36. Hodge, R. D. et al. Conserved cell types with divergent features in human versus mouse cortex. Nature 573, 61–68 (2019).

37. Tsanov, N. et al. smiFISH and FISH-quant – a flexible single RNA detection approach with super-resolution capability. Nucleic Acids Research 44, e165–e165 (2016).

38. Hershberg, E. A. et al. PaintSHOP enables the interactive design of transcriptome- and genome-scale oligonucleotide FISH experiments. Nature methods 18, 1265 (2021).

39. Moffitt, J. R. & Zhuang, X. RNA Imaging with Multiplexed Error-Robust Fluorescence In Situ Hybridization (MERFISH). in Methods in Enzymology vol. 572 1–49 (2016).

40. Aitken, C. E., Marshall, R. A. & Puglisi, J. D. An oxygen scavenging system for improvement of dye stability in sin gle-molecule fluorescence experiments. Biophysical Journal 94, 1826–1835 (2008).

41. Myronenko, A. & Xubo Song. Point Set Registration: Coherent Point Drift. IEEE Transactions on Pattern Analysis and Machine Intelligence 32, 2262–2275 (2010).

42. Rocklin, M. Dask : Parallel Computation with Blocked algorithms and Task Scheduling. Proceedings of the 14th Python in Science Conference 130–136 (2015).

43. Zhou, Q.-Y., Park, J. & Koltun, V. Open3D: A Modern Library for 3D Data Processing. (2018).

44. Hamilton, W. L., Ying, R. & Leskovec, J. Inductive Representation Learning on Large Graphs. Proceedings of the 31st International Conference on Neural Information Processing Systems 59 (2017).

